# Acquired epithelial Wnt7b secretion establishes autocrine WNT dependency in BRAF-mutant colorectal cancer

**DOI:** 10.64898/2026.09.24.754033

**Authors:** Simon R. Lampart, Tasnim Khadam-Al-Jame, Peter Leary, Ab Meijs, Clelia Pistoni, Tosca Dalessi, Randall J. Platt, Ralph Fritsch, Hassan Fazilaty, Tomas Valenta

## Abstract

Colorectal cancer (CRC) subtypes differ fundamentally in their reliance on WNT signaling, yet the sources and regulation of WNT ligands as well as the subtype-specific requirements for ligand-driven activation remain poorly defined. Here, we generated genetically-engineered organoid models of Apc- and Braf-mutant CRC to systematically dissect ligand dependency across distinct genetic backgrounds and CRC subtypes. Through genetic perturbation and pharmacological inhibition of WNT secretion, we find that BRAF-driven organoids critically depend on autocrine WNT ligand production for survival and proliferation. In contrast, APC-mutant organoids remain viable but undergo discernible transcriptional changes upon ligand withdrawal. We identify Wnt7b as a non-redundant epithelial WNT ligand required to sustain β-catenin–dependent signaling specifically in BRAF-mutant CRC. Chromatin accessibility and transcriptional profiling reveal that Wnt7b activation occurs early during transformation and coincides with cancer-specific opening of regulatory elements at the Wnt7b locus. Integrative regulatory analyses nominate the transcription factor RFX7 as a candidate upstream regulator of epithelial Wnt7b expression. Consistent with these findings, analysis of human CRC specimens demonstrates WNT7B expression in a subset of BRAF-mutant tumors and reveals an association with poor patient survival. Together, these results redefine ligand dependency in colorectal cancer and uncover autocrine WNT7B signaling as a mechanistically defined vulnerability in aggressive BRAF-mutant disease.

## Introduction

Colorectal cancer (CRC) progression is tightly coupled to deregulated WNT signaling^1–4^. In homeostasis, WNT signaling is essential for maintaining the colonic stem cell niche, with precise spatial and cellular control of ligand secretion^5,6^. Subepithelial mesenchymal populations, such as Gli1^+^/Pdgfra^high^ fibroblasts, are the main source of WNT ligands that sustain stem cell proliferation and control the WNT gradient along the crypt axis in the homeostatic colon^7,8^. In contrast, colon epithelial cells contribute only minimally to the WNT milieu in homeostasis^9,10^. During development and tissue repair however, epithelial WNT expression is transiently activated, demonstrating a context-dependent capacity of epithelial cells to re-engage WNT ligand production^11–13^. Emerging evidence suggests that tumor cells can also re-express WNT ligands in CRC, yet the functional significance, regulation and subtype specificity of this phenomenon is poorly described^14^.

Dysregulation of the WNT signaling pathway is a hallmark of CRC^1–3^, but the degree and mechanism of pathway activation vary substantially between the various molecular subtypes ^15–19^. APC-mutant tumors, which represent the majority of sporadic CRCs, activate WNT signaling downstream of the receptor^20,21^. Mutations in APC lock the pathway in a highly active state – irrespective of ligand binding^17,22,23^. By contrast, the BRAF-mutant CRC subtype follows a distinct route to malignancy^19,23,24^. It is associated with poor prognosis and WNT dysregulations at the receptor level ^4,20,25–27^. Frequent inactivation of the E3 ubiquitin ligases RNF43 and ZNRF3^4,27,28^ increases Frizzled (FZD) receptor abundance^29,30^ and sensitizes tumor cells to WNT ligands, suggesting a fundamentally different mode of pathway engagement. Accordingly, BRAF-mutant tumors are often classified as WNT-low ^17,20,23,31^. Whether this reflects strict ligand dependence, or instead a distinct regulatory architecture enabling alternative WNT activation, remains unresolved.

Emerging evidence has started to challenge the dichotomy between ligand-dependent and -independent WNT signaling. APC-driven cancer cells and organoids have been shown to retain responsiveness to exogenous WNT ligands, indicating residual ligand sensitivity^32,33^. Further, it was shown that WNT responses induced by receptor signaling differ from those induced by APC truncation^33^. Conversely, BRAF-mutant tumors frequently undergo transcriptional and epigenetic reprogramming that may enable WNT pathway activation independently of Wnt ligands^34–37^. These findings indicate that WNT dependency is more nuanced than previously recognized, highlighting the need to systematically define identity, source and regulation of WNT ligands across CRC subtypes. Several key questions remain unresolved. The genomic, epigenomic and transcriptomic mechanisms underlying epithelial WNT re-expression, and whether and whether tumor cells themselves establish autocrine WNT signaling programs, remain largely unknown. Progress in addressing these question has been constrained by the lack of physiologically relevant experimental models, particularly for BRAF-driven CRC.

To address this issue, we generated CRISPR-engineered CRC organoid models representing APC-mutant and BRAF-mutant subtypes. While both models survived independently of exogenous WNT ligands, BRAF-driven organoids uniquely relied on acquired autocrine WNT secretion. Through integrated transcriptomic and chromatin accessibility profiling, we examined the consequences of impaired WNT secretion in both subtypes, delineating subtype-specific differences in WNT dependency and exploring the regulatory basis of epithelial WNT re-expression. Moreover, we validated these findings in patient CRC samples.

Together, our models provide a framework to dissect subtype-specific WNT dependencies and illustrate how reactivation of epithelial WNT secretion shapes colorectal cancer biology.

## Results

### Establishment of Apc- and Braf-mutant colorectal cancer organoids displaying high and low WNT pathway activation

APC- and BRAF-driven CRC follow distinct routes to malignancy, whereas BRAF-mutant tumors are associated with significantly worse patient survival^23–26,38^ (**Fig.1a,b**). To explore the WNT dependency of these different CRC subtypes, we established two genetically defined, colonic organoid models recapitulating APC- and BRAF-mutant CRCs. To model APC-driven CRC, we sequentially introduced *Apc, KRAS^G12V^, Tp53* and *Smad4* mutations (AKPS) into murine wildtype (WT) organoids (**Fig.1c**), (**Fig.S1a,b,n**). This model is widely used to study aggressive WNT-driven colorectal cancer^39–41^. By withdrawal of the colonic niche factors WNT-surrogate, R-spondin 1 (Rspo), EGF and Noggin, we selected for cells harboring the desired genomic modifications and oncogenic MAPK activation was confirmed by pERK-immunoblotting (**Fig.1e**). The etiology of BRAF-driven CRC is less well-characterized. BRAF-driven tumors are frequently microsatellite instable (MSI) owing to epigenetic silencing of the mismatch repair protein MLH1^20,42^. BRAF-mutant, MSI tumors are associated with *RNF43* frameshift mutations due to coding microsatellite tracts in its sequence^43^. Its paralogue *ZNRF3* is more often inactivated by promoter hypermethylation^28^. For murine tumorigenesis, inactivation of both paralogues is necessary^18^. Similar to *RNF43*, also *TGFBR2* mutations are associated with microsatellite instability^44^. Accordingly, guided by prior reports that studied the mutational landscape of BRAF-mutant CRC^27,28,43–48^, we engineered prototypic organoids carrying *Braf^V637E^, Rnf43*, *Znrf3*, *Mlh1dn*, *Tgfbr2* and *Tp53* mutations (BMRZTP) to model this CRC subtype. (**Fig.1d**), (**Fig.S1c-n**). The activating *Braf^V637E^* mutation is the murine equivalent of human *BRAF^V600E^*which induces oncogenic MAPK signaling activation ^49^ (**Fig.1d**), (**Fig.S1g**). *Braf^V637E^*-mutant cells grew independent of EGF, were resistant to EGFR inhibition and showed increased phospho-ERK levels compared to Braf WT organoids (**Fig.1e**), (**Fig.S1g,h**). To mimic *MLH1* hypermethylation, we overexpressed a dominant negative *Mlh1* allele (*Mlh1dn*), which suppresses the function of wildtype *Mlh1*^50^. Fragment length analysis of five validated murine MSI markers revealed instability at 2/5 loci, consistent with MSI-H status in analogy to the human Bethesda panel^51,52^ (**Fig.1f**), (**Fig.S1i,j**). Removal of the RING domain required for FZD ubiquitination in *Rnf43* and *Znrf3,* allowed outgrowth of mutant, but not WT organoids upon removal of Rspo from the culture medium (**Fig.S1c-e**). *Tgfbr2*-mutant cells were selected by TGFβ treatment and Noggin withdrawal (**Fig.S1k**). Finally, to model more aggressive stages, we also targeted *Tp53* and selected mutant cells using Nutlin-3a (**Fig.S1l**).

**Figure 1.**
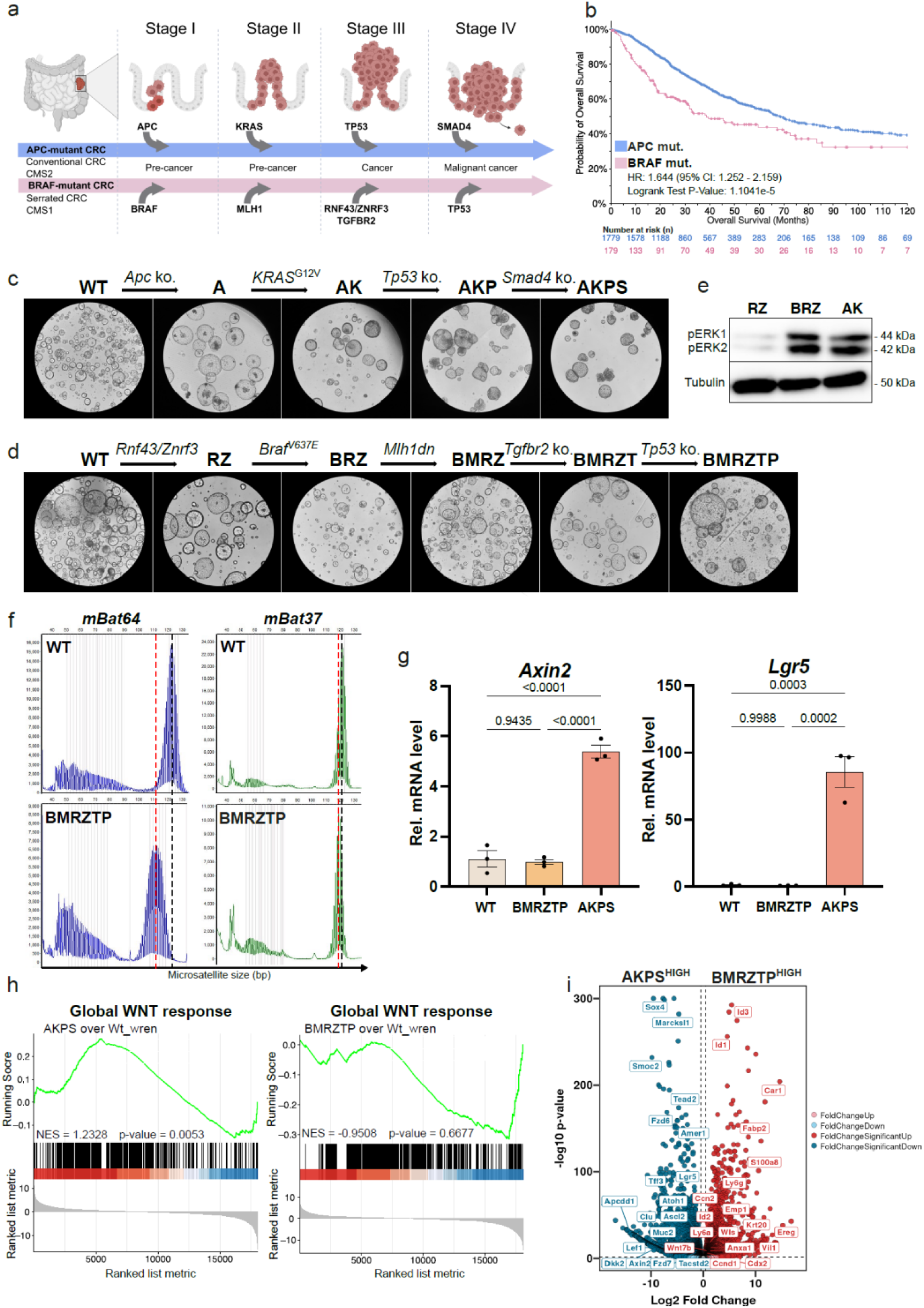
Development of High- and low-WNT Apc- and Braf-mutant CRC organoids. a) Schematic depiction of APC- and BRAF-mutant CRC progression starting from benign precursor lesions to malignant carcinomas. CMS = Consensus molecular subtype. Scheme was created with BioRender.com. b) Overall survival of CRC patients with APC or BRAF mutations over a time period of 120 months. Kaplan-Meier curve was generated using the cBioportal. Hazard ratio and Logrank test p-value were calculated based on 1779 patients with APC and 179 patients with BRAF mutation. c) Stepwise generation of *Apc* ko., lenti-*KRAS*^G12V^, *Tp53* ko., *Smad4* ko. (AKPS) organoids starting from wild type (WT) Villin-Cre^ERT2^; *Wls*^fl/fl^ organoids. d) Stepwise generation of lenti-*Braf*^V637E^, *Rnf43* ΔRING domain (RD), *Znrf3* ΔRD, *Mlh1* dominant negative (*Mlh1*dn), *Tgfbr2* ko., *Tp53* ko. (BMRZTP) organoids starting from WT Villin-Cre^ERT2^; *Wls*^fl/fl^ organoids. *Braf*^V637E^ is the murine equivalent to the human *BRAF*^V600E^ allele. e) Activation of the MAPK pathway visualized by phospho-ERK immunoblot. f) Microsatellite fragment length analysis (FLA) of healthy WT and cancerous BMRZTP organoids. The two loci mBat64 and mBat37 are shown. Dashed black line indicates average length in WT organoids. Red dashed line indicates average length in BMRZTP organoids. g) Relative mRNA levels of the WNT target genes *Axin2* and *Lgr5* in organoids cultured 48 h without WNT surrogate and R-spondin 1. P-value was calculated by ordinary one-way ANOVA using Tukey’s multiple comparison test. h) Gene set enrichment analysis (GSEA) of the Global_WNT response signature from Kleeman et al. 2020^53^. Analysis was performed using bulk RNA sequencing data of BMRZTP, AKPS and WT organoids (Wt_wren) starved of all niche factors. Gene sets are displayed as running score plot. Green line shows enrichment of the gene sets in AKPS and BMRZTP over Wt_wren. Vertical black bars indicate the position of genes within the ranked list. Normalized enrichment score (NES) and p-value are shown. i) Volcanoplot displaying differential gene expression between BMRZTP and AKPS organoids. Threshold for significance was set at log2FC > 0.5, FDR < 0.05. Selected genes are labelled.

To measure intrinsic WNT activation, organoids were cultured in medium lacking WNT surrogate (W) and Rspo (R) followed by interrogation of the WNT target genes *Lgr5* and *Axin2* (**Fig.1g**). Compared to BMRZTP and WT, AKPS organoids showed high *Axin2* and *Lgr5* levels, also in the absence of WNT surrogate. In contrast, both BMRZTP and WT organoids displayed comparably low WNT activation. Bulk RNA sequencing followed by gene set enrichment analysis (GSEA) revealed an enrichment of the Global_WNT response signature^53^ in AKPS but not BMRZTP (**Fig.1h**). Similarly, compared to AKPS, BMRZTP lacks a significant activation of several additional WNT target genes (*Lef1*, *Ascl2*, *Apcdd1*, *Amer1*, *Dkk2*) (**Fig.1i**).

Taken together, these models capture the key genetic features of APC- and BRAF-driven CRC and mirror the high- and low-WNT activation observed in patients^17,19,20,23^. They provide a genetically defined platform to explore how subtype-specific molecular programs influence colorectal cancer.

### Autocrine WNT secretion drives survival and proliferation of BRAF-mutant but not APC mutant CRC organoids

We next investigated how WNT ligands contribute to organoid survival and growth. We first compared the growth of AKPS and BMRZTP organoids in −WR medium (lacking WNT surrogate, Rspo, EGF and Noggin). Additionally, BMRZT organoids were included to monitor *Tp53*-independent behavior, as Tp53 was previously implicated in conferring niche independency^54^. The growth of AKPS organoids was not affected by WR withdrawal (**Fig.2a**), however, to our surprise, also BMRZT and BMRZTP organoids could be passaged continuously in medium devoid of all niche factors (**Fig.2a**). This is in stark contrast to WT organoids which did not survive passaging without exogenous WNT surrogate and Rspo1 (−WR), even in the presence of EGF and Noggin (**Fig.S1a**). This observation suggested that cell-intrinsic WNT secretion could act as an alternative WNT source.

**Figure 2.**
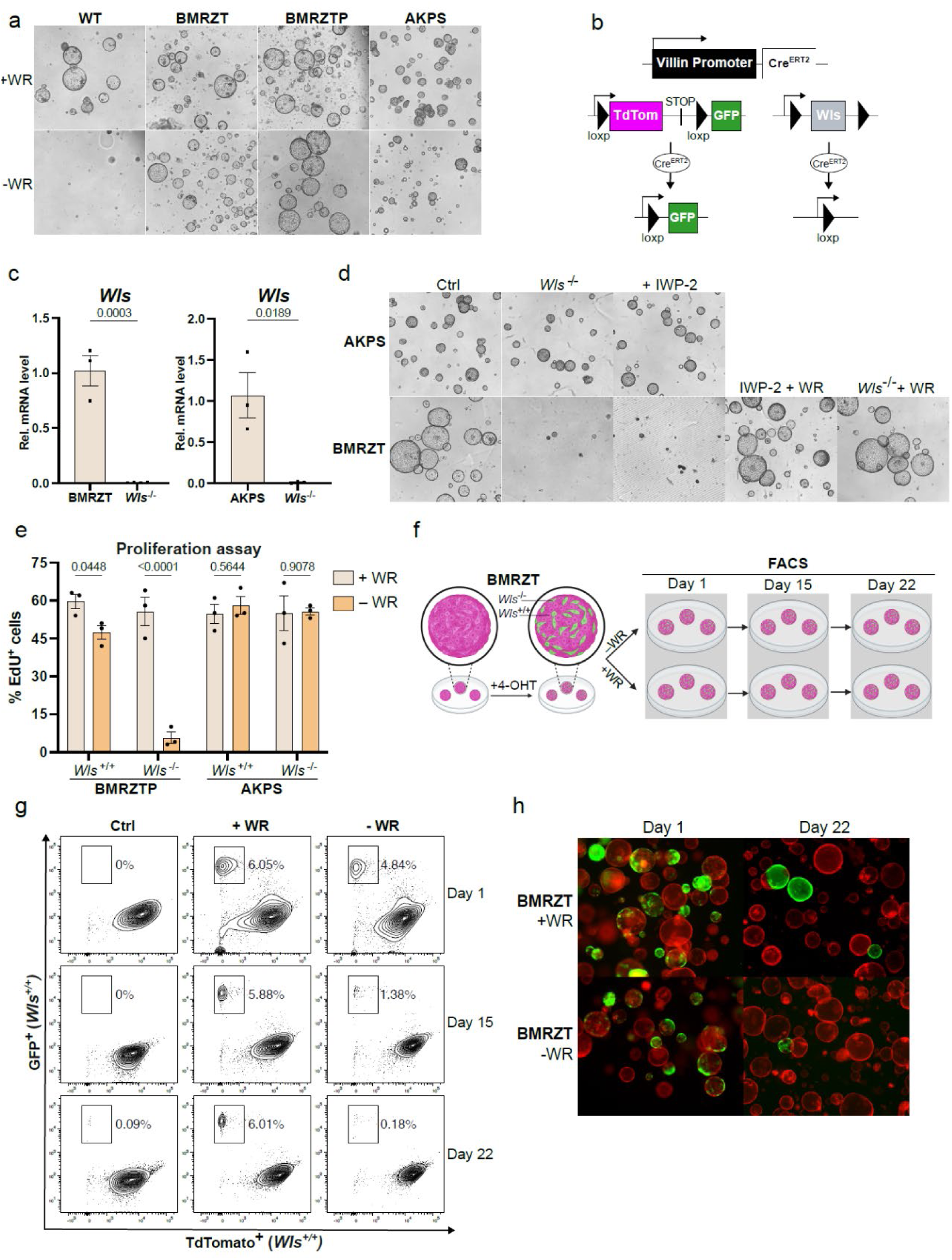
BMRZT and BMRZTP but not AKPS organoids are dependent on autocrine WNT secretion. a) WNT starvation of WT, BMRZT, BMRZTP and AKPS organoids. AKPS and Braf-derived organoids were either cultured with WNT surrogate and R-spondin 1 (+WR) or without all niche factors (WNT surrogate, R-spondin 1, EGF and Noggin) (−WR). For WT organoids, EGF and Noggin were added additionally in both conditions. Representative images after two passages in the respective media are shown. b) Schematic overview representing the *R26*-*mTmG*; *Villin*-_Cre_^ERT2^; *Wls*^fl/fl^ background present in all organoids used in this study. The *mTmG* and the *Villin*-Cre^ERT2^ loci are expressed from different chromosomes. At baseline, organoids express TdTomato and are *Wls* proficient. Upon Cre induction by 4-hydroxy Tamoxifen (4-OHT), the *Wls* and *mTmG* loci are independently recombined resulting in GFP^+^; *Wls*^−/−^organoids. c) Relative *Wls* mRNA levels in BMRZT and AKPS before and after *Wls*^−/−^recombination and FACS sorting of GFP^+^ cells. P-value for both was calculated using an unpaired t-test. d) Testing the effect of WNT secretion blockade in medium –WR. WNT secretion was either blocked by *Wls* knock-out or treatment with the Porcupine inhibitor IWP-2. Phenotypes were rescued by concomitant WR supplementation. Representative images after two passages are shown. e) EdU proliferation assay of recombined or unrecombined BMRZTP or AKPS organoids. Organoids were cultured for one passage in +WR or −WR medium. % of proliferating EdU^+^ cells are shown. n = 3. P-values were calculated using ordinary two-way Anova with Sidak’s multiple comparison test. f) Schematic overview of the time course experiment. Chimeric BMRZT organoids consisting of *Wls*^+/+^ / *Wls*^−/−^cells were generated by addition of 4-OHT. Organoids were then divided into a −WR and a + WR condition and the percentage of live GFP^+^(*Wls*^−/−^) cells was assessed by FACs after 1, 15 and 22 days in the respective medium. Created with BioRender.com) FACS dot plots including the gate used to measure the percentage of live BMRZT GFP^+^ cells in the +WR and −WR condition. Timepoint of acquisition is indicated on the right side. Unrecombined BMRZT organoids were used as control. h) Representative images of chimeric BMRZT organoids in the + WR and −WREN condition at day 1 and day 22 of the experiment. TdTomato (magenta) marks unrecombined cells, GFP (green) marks recombined cells.

To test our hypothesis, we blocked endogenous WNT secretion by genetic removal of *Wls,* which is the essential transporter required for the secretion of all WNTs^55^ or the addition of a Porcn inhibitor (Porcni), IWP-2, which inhibits WNT palmitoylation necessary for secretion^56^ (**Fig.2c,d**), (**Fig.S2b,c**). Given the background of our organoids (*R26*-*mTmG*; *Villin*-Cre^ERT2^; *Wls*^fl/fl^), *Wls* recombination coincided with a fluorophore switch from TdTomato to GFP (**Fig.2b**), (**Fig.S2a**). AKPS *Wls*^−/−^organoids showed no visible phenotype upon WR withdrawal or IWP-2 treatment (**Fig.2d**), (**Fig.S2c**). Likewise, proliferation rates between AKPS and AKPS *Wls*^−/−^were comparable in both WR-containing and WR-depleted medium. (**Fig.2e**), (**Fig.S2d**). Conversely, both BMRZT *Wls*^−/−^ and BMRZTP *Wls*^−/−^organoids did not survive passaging, which was preceded by a loss of proliferative cells (**Fig.2d,e**), (**Fig.S2c,d**). This phenotype was recapitulated by IWP-2 treatment and could be rescued by WR supplementation.

These results demonstrate that BRAF-driven organoids depend on endogenous secretion of WNT ligands for survival and proliferation, while APC-driven organoids proliferate independent of WNT secretion. WNT ligands are highly lipophilic molecules and their range of action is debated ^6,57,58^. Thus, we tested, if short-range paracrine secretion is sufficient to sustain survival and proliferation in neighboring cells. Using 4-Hydroxytamoxifen (4-OHT) treatment, we generated chimeric BMRZT organoids composed of *Wls*-proficient and *Wls*-deficient cells expressing TdTomato or GFP, respectively. Subsequently, organoids were cultured with or without WR and monitored over the course of three weeks (**Fig.2f-h**), (**Fig.S2e**). Over time, the GFP^+^ population drastically decreased in medium lacking WR while it was stable in +WR medium. This indicates that paracrine WNT secretion is insufficient to drive survival and proliferation of neighboring cells, suggesting autocrine secretion to be the primary WNT source.

In summary, we show that Braf-mutant but not Apc-mutant organoids are dependent on autocrine WNT secretion for survival and proliferation, a dependency that is unaffected by Tp53 mutational status. These results reveal a previously unappreciated autocrine loop mechanism sustaining basal WNT signaling in Braf-mutant CRC.

### Loss of WNT secretion drives differentiation, cellular stress and apoptosis in WNT-dependent CRC organoids

Following the determination of WNT dependency and independency in our models, we next determined the consequences associated with the loss of autocrine secretion. To test this, we performed bulk RNA sequencing of AKPS *Wls^−/−^* and BMRZTP *Wls^−/−^*cultured in medium lacking WR. Principal component analysis (PCA) revealed tight clustering of AKPS and AKPS *Wls^−/−^*organoids, indicating a limited transcriptomic response to *Wls* recombination (**Fig.3a**). By contrast, consistent with the effect on survival, both BMRZT *Wls^−/−^* and BMRZTP *Wls^−/−^*differed substantially from their unrecombined counterparts. Intriguingly, BMRZT *Wls^−/−^* and BMRZTP *Wls^−/−^*were transcriptionally similar. These results are aligned with the previously observed phenotype and suggest that the effects of *Wls* loss are independent of *Tp53* mutation. IWP-2-mediated Porcn inhibition (BMRZT_Porcni) mirrored *Wls* loss, as reflected by the overlapping differentially expressed genes (DEGSs) between BMRZT *Wls^−/−^* and BMRZT_Porcni organoids (**Fig.3b**), (**Fig.S3j**).

**Figure 3.**
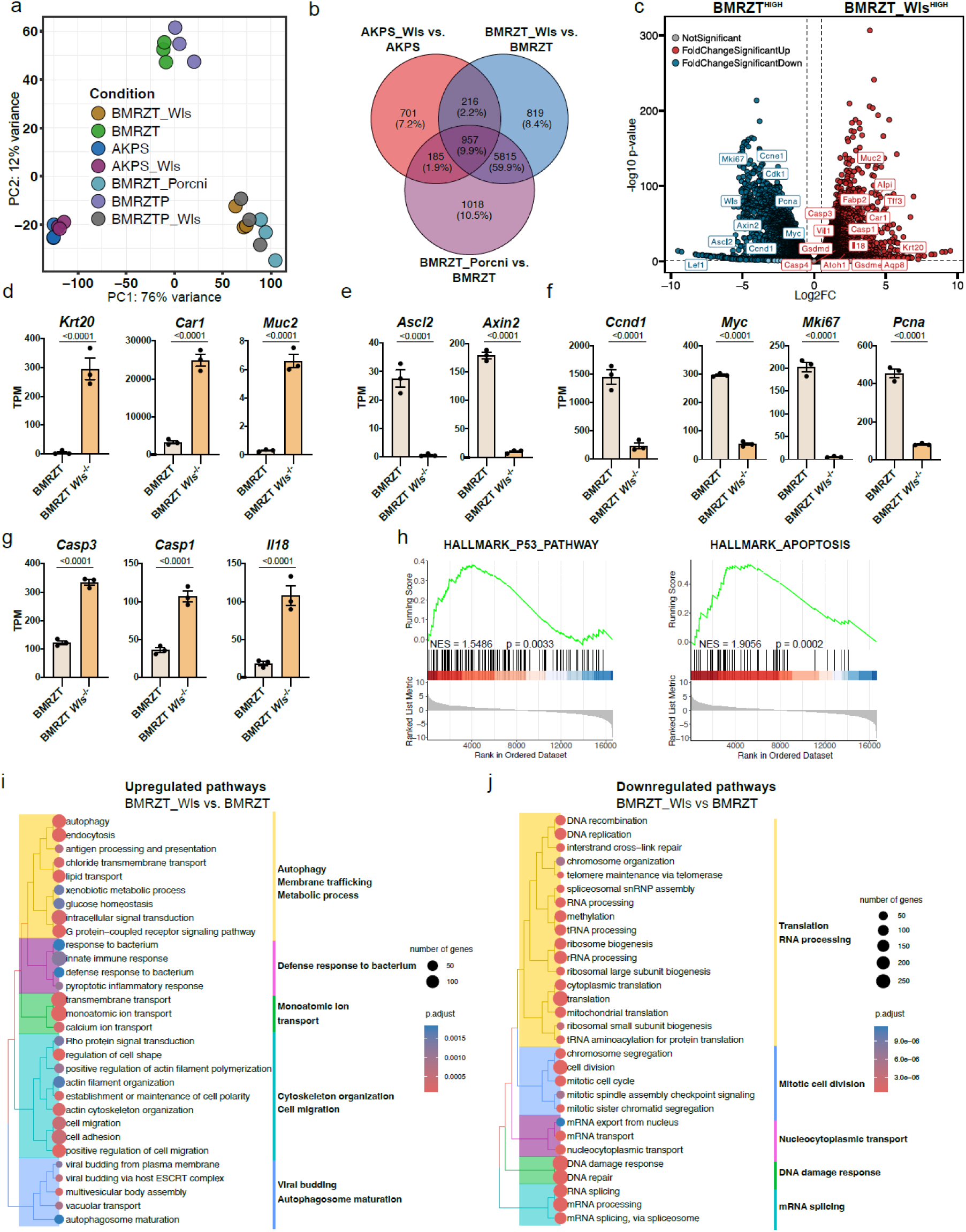
Loss of WNT secretion in BMRZT induces differentiation and apoptosis. a) Principal component analysis (PCA) plot of bulkRNAseq datasets. Each point represents one sample, colored by condition. Percentage of variance explained is indicated on the axes. b) Venn diagram displaying the overlap between differentially expressed genes (DEG) upon loss of WNT secretion in BMRZT and AKPS organoids. WNT secretion was blocked either by *Wls* recombination (BMRZT_Wls) or by Porcupine inhibition via IWP-2 (BMRZT_Porcni). In both conditions, RNA was collected 48 h after transferring organoids to medium lacking all niche factors and start of IWP-2 treatment. Up and downregulated DEGs with log2FC > 0.5 and FDR < 0.05 are shown. c) Volcano plot showcasing transcriptomic differences between BMRZT_Wls (lacking *Wls*) and BMRZT organoids. Threshold for significance was set at log2FC > 0.5, FDR < 0.05. Selected genes are labelled. d) mRNA expression levels of differentiation markers in the bulk dataset between BMRZT and BMRZT_Wls. mRNA counts are shown as transcript per million (TPM). e) Expression levels of WNT target genes. f) Expression levels of proliferation associated genes. g) Expression levels of apoptosis (*Casp3*) and pyroptosis markers (*Casp1*, *Il18*). FDR values in d), e), f) and g) were calculated by DESeq2 as part of the bulk mRNA sequencing analysis. h) Gene set enrichment analysis (GSEA) on the HALLMARK_P53 and HALLMARK_APOPTOSIS gene sets displayed as running score plots. Green line shows enrichment of the genesets in BMRZT_Wls over BMRZT. Vertical black bars indicate the position of genes within the ranked list. Normalized enrichment score (NES) and p-value are shown. i-j) Over representation (ORA) analysis of the top 30 up- and downregulated pathways between BMRZT_Wls and BMRZT summarized in tree-plots. The ORA analysis included all genes with a raw p-value < 0.01 and log2FC < 0.5. Results are based on the “Molecular function” gene ontology (GO). Number of genes per GO term are indicated by dot size. Pathway summary was added manually. Color indicates statistical significance of pathway enrichment. Adjusted p-values are shown.

Close examination of differentially expressed genes (DEGs) between BMRZT *Wls^−/−^* and BMRZT revealed an upregulation of differentiation markers (**Fig.3c,d**). In particular, mature enterocyte markers (*Vil1*, *Krt20*, *Car1*, *Aqp8*, *Fabp2*), but also goblet cell markers (*Atoh1*, *Muc2*, *Tff3*) were significantly elevated. Moreover, we detected an increase in genes associated with apoptosis and pyroptosis (*Casp1*, *Casp3*, *Gsdme*, *Il18*) (**Fig.3c,g**). This was accompanied by an enrichment of apoptotic signaling signatures (**Fig.3h**). Among the downregulated genes, we found factors involved in proliferation and cell cycle progression (*Myc*, *Pcna*, *Mki67 Cdk1*, Ccnd1, Ccne1), but also WNT target genes (*Axin2*, *Ascl2*, *Lef1*) (**Fig.3c,e,f**).

Global pathway analysis identified an upregulation of autophagy, lipid and ion transport, implying increased recycling of organelles, nutrient flux and membrane trafficking (**Fig.3i**). These changes coincided with increased cytoskeletal remodeling and pathways involved in cell adhesion. In parallel, metabolic rewiring and activation of inflammatory and innate immune pathways were indicative of a cellular stress response. Analysis of downregulated pathways revealed that both cytoplasmic and mitochondrial translation were heavily decreased, suggesting that *Wls* loss leads to a strong impairment of the protein synthesis machinery (**Fig.3j**). In similar fashion also RNA processing, maturation and transport pathways were reduced, likely exacerbating translational stress. Further, programs responsible for DNA replication, repair and cell cycle control were significantly downregulated, suggesting replication stress and defective cell division.

Even though we previously demonstrated that AKPS organoids displayed no significant changes in morphology or proliferation following blockage of WNT secretion, we still detected discernible transcriptomic differences between AKPS *Wls*^−/−^ and AKPS organoids (**Fig.S3a,g**). Despite their constitutive WNT activation, we saw a trend towards downregulation of WNT target genes (*Axin2*, *Notum*, *Apcdd1*) and also canonical stem cell markers (*Lgr5*, *Ascl2*) (**Fig.S3b,c**). On the other hand, we detected an upregulation of fetal markers (*Anxa1*, *Ly6a*, *Tacstd2*), Yap targets (*Ccn1*, *Ccn2*) (**SFig3e**) and *Emp1* which marks invasive *Lgr5*-cells^59^ (**Fig.S3d,e,f**). Among the enriched pathways, we found an upregulation of cytoskeletal remodeling, migratory and angiogenic programs, highlighting a shift towards structural adaptation and tissue remodeling (**Fig.S3h**). Concomitantly, we detected a downregulation of cell adhesion, and pathways important during development and morphogenic patterning (**Fig.S3i**).

Taken together, loss of autocrine WNT secretion in BRAF-driven organoids leads to terminal differentiation and an adaptive stress response involving autophagy and inflammatory programs. In addition, downregulation of the translation and cell cycle machinery suggest a strong impairment of biosynthetic and proliferative processes. Given that BMRZT *Wls^−/−^*organoids do not survive passaging in medium lacking WR, these processes likely proceed apoptotic cell death. AKPS organoids, albeit WNT independent, remain responsive to lack of WNT ligands and display subtle transcriptional changes that go beyond downregulation of WNT target genes.

### Autocrine Wnt7b is essential for survival and growth in BMRZTP organoids by driving the WNT/β-catenin pathway

Since autocrine WNT secretion was indispensable for the survival of BRAF-driven organoids, we next aimed to delineate the WNT code, i.e. the spectrum of WNT ligands orchestrating these pro-survival effects. Strikingly, among all 19 WNTs, Braf-mutant models expressed only *Wnt7b* while WT organoids had negligible WNT expression (**Fig.4a,b**), (**Fig.S4a**). In AKPS organoids, *Wnt7b,* besides *Wnt3*, *Wnt5b* and *Wnt6,* was the highest expressed WNT (**Fig.S4a,b**). These findings demonstrate, that epithelial WNT expression is conserved between CRC subtypes harboring different WNT activation.

**Figure 4.**
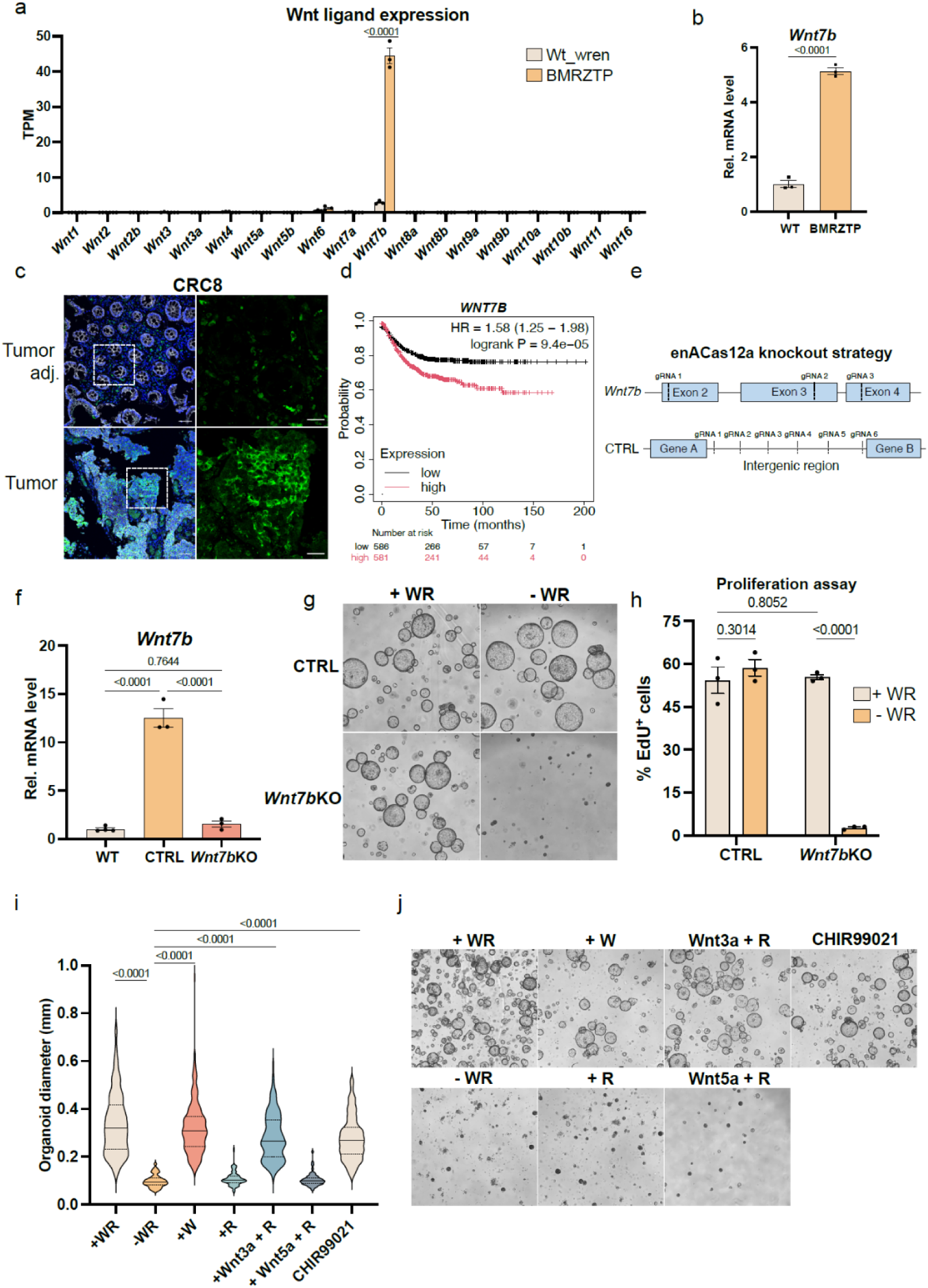
BMRZTP organoids are dependent on autocrine Wnt7b secretion. a) Bar chart displaying mRNA counts in transcripts per million (TPM) of all 19 WNT ligands in BMRZTP and WT (WT_wren) organoids. FDR value was calculated by DESeq2. b) Validation of *Wnt7b* expression by RT-qPCR. P-value was calculated using an unpaired t-test. c) IF staining on human BRAF-mutant tumor and matched tumor-adjacent tissue. Staining shows WNT7B (green), E-cadherin (white) and DAPI (blue). Dashed white boxes indicate insets shown in right. Scalebars: 60 and 25 μm. d) Kaplan-Meier plots showing the inverse correlation between *WNT7B* expression and CRC patient relapse free survival. Hazard ratio (HR) and logarithmic ranked p-value (logrank P) was calculated between high (red) and low (black) expressing cohorts. Numbers below indicate number of patients at risk over time (months). Statistical analysis was performed on the default settings. e) Schematic overview of the enACas12a knockout strategy to knock out *Wnt7b* in BMRZTP organoids. For *Wnt7b*, three exons were targeted using three gRNAs. For CTRL organoids, an intergenic region was targeted with six gRNAs. f) Loss of *Wnt7b* transcript levels in *Wnt7b*KO compared to WT and CTRL organoids. P-value was calculated by ordinary one-way ANOVA using Tukey’s multiple comparison test. g) WNT starvation of CTRL and *Wnt7b*KO organoids cultured in either +WR or −WR medium. Images shown were taken after two passages in the respective media. h) EdU proliferation assay of CTRL and *Wnt7b*KO organoids cultured either in +WR or −WR medium. Percentage of EdU^+^ proliferating cells is shown. P-values were calculated using two-way Anova with Fisher’s LSD test. i) Rescue of *Wnt7b*KO organoids. Violin plot depicts diameters of *Wnt7b*KO organoids grown in the different media for two passages. Plots show pooled diameters of three independent experiments. Per replicate and condition at least 25 organoids were measured. P-values were calculated based on the mean organoid length per replicate and condition by ordinary one-way Anova with Tuckey’s multiple comparison test. j) Representative images of *Wnt7b*KO organoids taken 48h after the second passage. To activate canonical WNT signaling, WNT surrogate and Rspo (+WR), WNT surrogate only (+W), CHIR99021 (GSK-3 inhibitor) and recombinant Wnt3a + Rspo were used. Non-canonical WNT signaling was induced by recombinant Wnt5a + Rspondin1 supplementation. The +Rspo (+R) and −WR conditions served as negative control.

As Wnt7b-mediated pathway activation depends on the availability of matching Fzd receptors, we next examined the Fzd profile present in both models (**Fig.S4c**). In AKPS, *Fzd6* and *Fzd7* showed the highest expression compared to BMRZT and BMRZTP in which *Fzd1* and *Fzd5* –similar to WT–were the most prominent. Co-receptors *Lrp5* and *Lrp6* were increased in both AKPS and BMRZTP compared to WT (**Fig.S4d**). As Wnt7b was previously shown to bind to Fzd1, Fzd5, Fzd7 and Fzd10^60–64^ both models have the potential to engage canonical (i.e. β-catenin-dependent) Wnt7b-Fzd signaling through one or more of these receptors.

To assess the clinical relevance of our findings, we next investigated if *WNT7B* expression is also a feature of human CRC. Immunostaining on *BRAF*-mutant CRC patient samples revealed robust WNT7B expression in one out of three patients (**Fig.4c**), (**Fig.S4e**). Expression levels in matching tumor-adjacent tissue were negligible. Remarkably, high expression of *WNT7B* in human CRC correlated with inferior patient prognosis (**Fig.4d**) which further highlights the clinical relevance of our finding and its role in tumor formation and progression.

To directly study the role of Wnt7b, we generated *Wnt7b* knockout (*Wnt7b*KO) BMRZTP organoids. By employing enhanced Cas12a (enCAs12a)-mediated genome editing, we targeted the *Wnt7b* coding sequence in three different exons (**Fig.4e,f**) (**Fig.S4f,g**). Simultaneously, we generated control (CTRL) BMRZTP organoids using the same method and sgRNAs targeting non-coding intergenic regions. Next, to test specific dependency on autocrine Wnt7b secretion, we cultured *Wnt7b*KO and CTRL organoids in medium lacking WR. *Wnt7b*KO organoids did not survive passaging in −WR medium compared to CTRL organoids (**Fig.4g**) Further, cell death in *Wnt7b*KO was preceded by a significant drop in proliferation compared to CTRL organoids (**Fig.4h**), (**Fig.S4h**). Notably, this phenotype was rescued by WR supplementation which restored proliferation levels to that of CTRL organoids.

Wnt7b has been shown to activate both β-catenin-dependent (canonical) and independent WNT signaling^65–69^. To evaluate by which branch its effects are governed, we attempted to rescue *Wnt7b*KO organoids using different supplements activating both pathways (**Fig.4i,j**). β-catenin-dependent pathway activation by supplementation with WNT surrogate, recombinant Wnt3a + Rspo1 or the GSK3-inhibitor CHIR99021, but not Rspo alone, rescued organoid growth. β-catenin independent activation by Wnt5a + Rspo1 also did not rescue the phenotype (**Fig.4i,j**).

In conclusion, we show that BMRZTP organoids are specifically dependent on autocrine Wnt7b secretion which promotes survival and proliferation by activating canonical, β-catenin-dependent WNT signaling.

### *Wnt7b* expression in precancerous cells is linked to differential chromatin accessibility

Having shown that *Wnt7b* is expressed in both murine and human CRC and plays a crucial role in *Braf*-driven organoids, we set out to study the mechanisms regulating the induction of *Wnt7b* expression. As both CRC models were generated in a stepwise fashion, we tested at which stage *Wnt7b* expression appears and which mutations coincide with its onset. To our surprise, we already detected *Wnt7b* expression in our earliest models consisting of *Apc*-mutant (A) and *Rnf43*/*Znrf3*-mutant (RZ) organoids (**Fig.5a**). Akin to BMRZT, BMRZTP and AKPS, *Wnt7b* was the only WNT ligand expressed in RZ and A organoids (**Fig.S5b**). RZ organoids also displayed a strong increase in *Wls* expression compared to WT organoids (**Fig.S5e**). Based on these results, we assessed, whether RZ organoids could survive in the absence of WR. Remarkably, they remained viable, demonstrating that autocrine *Wnt7b* signaling is established already in early precancerous stages (**Fig.5b**). Culture of RZ organoids in full medium or without WR did not significantly change *Wnt7b* levels (**Fig.S5c**). Also WT organoids in −WR medium did not upregulate *Wnt7b* (**Fig.S5d**). This suggests that neither high nor low WNT levels are likely to be direct triggers of *Wnt7b* expression.

**Figure 5.**
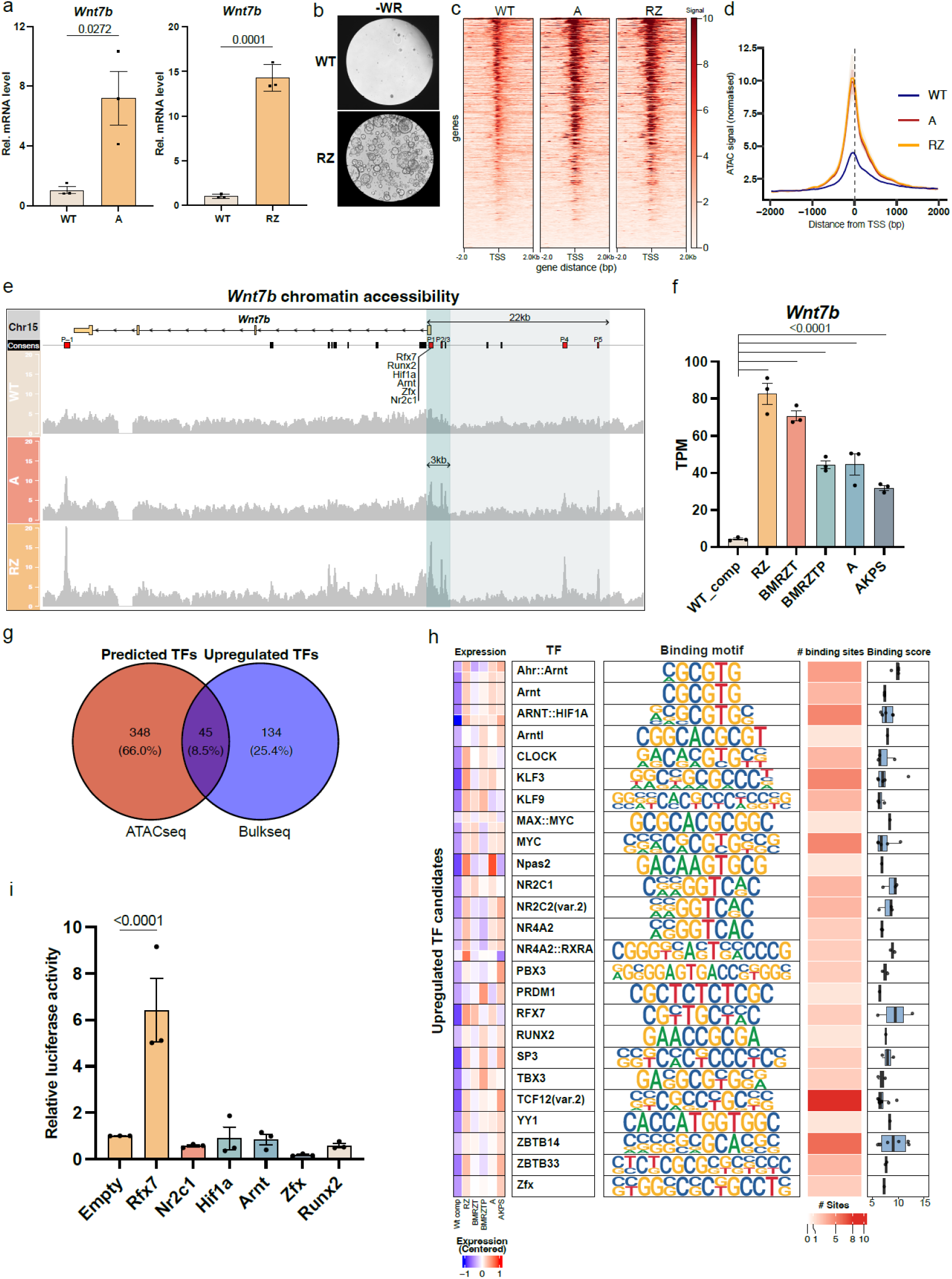
*Wnt7b* expression arises early during tumorigenesis and is associated with differential chromatin accessibility. a) RT-qPCR showing *Wnt7b* expression in the pre-cancerous *Rnf43*/*Znrf3* mut. (RZ) and *Apc* ko. (A) organoids. P-value in both was calculated via unpaired t-test. b) Representative images showing survival of RZ but not WT organoids in −WR medium supplemented with EGF and Noggin. Images were taken after two passages. c) Heatmap of chromatin accessibility around the transcriptional start site (TSS) in WT_comp, A and RZ organoids. ATAC-seq signal profile +/−2kb around the TSS is shown. d) Quantification of the normalized ATAC signal around the TSS in Wt_comp (WT organoids in full medium), A and RZ organoids. e) Overview of the ATAC-seq peak profile in WT, A RZ organoids around the *Wnt7b* locus on murine chromosome 15. Yellow boxes represent the four *Wnt7b* exons on the anti-sense strand, transcribed from right to left. Boxes below represent the peaks present in the consensus peak set based on all 3 conditions. Red boxes indicate the candidate peaks included into the TF binding site analysis. P-1 marks a 3’ downstream peak. Candidate TFs binding to the peak closest to the TSS (P1) are shown below P1. Cyan colored box indicates the proximal promoter region of 3kb length. Grey box indicates the extended regulatory region 22kb upstream of the TSS. f) Comparison of *Wnt7b* levels in the bulkRNAseq dataset in *Wnt7b*-expressing conditions compared to WT (Wt_comp). FDR values were calculated by DESeq2. TPM values of RZ, BMRZT, BMRZTP, A and AKPS are also shown in Fig.4a, Fig.S4a, Fig.S5b. g) Venn diagram showing the overlap between TFs with predicted binding to any of the candidate peaks (P-1 – P5) with binding score > 6 and TFs upregulated in RZ, BMRZT, BMRZTP, A and AKPS compared to Wt_comp). (log2FC > 0, FDR < 0.05). h) Table displaying the binding motifs and number of binding sites of the 22 significantly upregulated TFs with predicted binding to P1. Expression values are derived from bulkRNAseq. Binding scores were calculated using the monaLisa package. i) Relative luciferase activity in HEK293T cells measured upon transfection of candidate TFs binding to the proximal promoter region of *Wnt7b* (3bk including P1, P2 and P3). P-value was calculated by one-way ANOVA with Dunnet’s multiple comparison test.

RZ and A organoids both harbor mutations affecting the same pathway, yet correspond to distinct tumorigenic trajectories. The parallel expression of *Wnt7b* in both contexts prompted us to explore, whether shared regulatory mechanisms underlie its activation. To examine changes in chromatin accessibility associated with *Wnt7b* expression, we performed Assay for transposase accessible chromatin sequencing (ATAC-seq) of RZ, A and WT organoids (**Fig.5c-e**), (**Fig.S5g,f**). Analysis of transcriptional start sites (TSS) uncovered a striking change of the chromatin landscape in RZ and A organoids (**Fig.5c**). Compared to WT, accessibility of TSS in RZ and A was markedly increased (**Fig.5d**). More detailed examination of the *Wnt7b* locus exposed multiple ATAC peaks (P1 – P5) in RZ and A indicating accessible chromatin in the 22kb region including the cis-regulatory upstream sequence of the *Wnt7b* TSS (**Fig.5e**). In addition, we also detected a strong peak in a *Wnt7b*-adjacent downstream enhancer region (P-1). To discern putative transcription factors (TFs) regulating *Wnt7b* expression, we cross-referenced DEG TFs in all *Wnt7b*-expressing organoids (A, RZ, BMRZT, BMRZTP and AKPS) compared to WT, with TFs predicted to bind to ATAC-peak regions P-1 – P5, (**Fig.5 f,g**). Out of the 179 upregulated TFs, 45 displayed predicted binding, 22 of which had at least one binding site in P1, the peak closest to the TSS (**Fig.5h**). Among the 22 transcription factors analyzed, Rfx7, Runx2, Hif1a, Arnt, Zfx, Nr2c1 were selected as candidates regulating *Wnt7b,* based on their expression levels and binding scores (**Fig.5e**), (**Fig.S5h,i**). The expression of *Rfx7* and *Nr2c1* showed significant correlation with *Wnt7b* mRNA levels across all *Wnt7b*-expressing samples (**Fig.S5i**).

To validate DNA binding of these candidates, we performed luciferase reporter assays by overexpressing candidate TFs in HEK293T cells together with a reporter construct harboring the proximal, 3kb *Wnt7b* promoter region (**Fig.5i**). Here, Rfx7 displayed robust binding to the *Wnt7b* promoter region making it a top candidate controlling Wnt7b expression.

In conclusion, our experiments show that *Wnt7b* expression arises already in precancerous stages of both Apc- and Braf-mutant CRC and that its expression is controlled by shared programs across genomic backgrounds with Rfx7 as a likely direct regulator of expression.

## Discussion

In this study, we established two novel organoid models recapitulating *Apc*-mutant and *Braf*-mutant CRC to dissect subtype-specific WNT dependencies. While AKPS organoids were independent of WNT secretion for growth, they remained responsive to its ablation, whereas BMRZT/BMRZTP organoids critically depended on autocrine secretion of Wnt7b. Both subtypes show *de-novo* expression of *Wnt7b,* emerging in precancerous but not healthy organoids, accompanied by differential chromatin accessibility and shared transcriptional regulatory programs. Collectively, we uncover Wnt7b as a context-specific effector and potential therapeutic target in BRAF-driven CRC.

BRAF-mutant CRC represents a rare and aggressive subtype characterized by poor clinical outcome, especially following relapse^25,26,38^. While several studies have sought to model this disease^47^, an experimental framework analogous to the AKPS model for APC-mutant CRC has been lacking. Our BMRZTP model recapitulates key molecular and phenotypic features of BRAF-driven patient tumors, including their low-WNT background, and thus helps to address this gap. This model enables systematic investigation of BRAF-mutant CRC biology and the implications of WNT secretion in WNT-dependent CRC.

BRAF-driven CRC with RNF43 mutations is often described as WNT hypersensitive but still dependent on an exogenous WNT source^18^. Here, we show that upregulation of autocrine WNT secretion can confer complete independence from exogenous WNT ligands and R-spondin. While we are among the first to report complete WNT self-sufficiency in a murine model of BRAF-driven CRC^28^, this phenomenon also occurs in human CRC. Several studies found that BRAF-mutant patient-derived CRC organoids (PDOs) are able to survive without WNT supplementation and in some cases even in the absence of both WNT and R-spondin ^34,70–73^. These organoids displayed PORCN inhibitor sensitivity, consistent with a reliance on autocrine WNT secretion. Notably, the reported levels of WNT self-sufficiency differs across studies. An explanation for this may be varying mutational background, especially distinct RNF43 mutations, which are known to produce variable phenotypes^74,75^.

In stark contrast to BMRZTP, growth and proliferation in AKPS was completely independent of WNT secretion. Nevertheless, AKPS organoids remained responsive to the loss of WNT secretion by a downregulation of stem and WNT target genes and a concomitant modest increase in fetal markers, such as Tacstd2. This is consistent with previous studies showing that cells harboring truncated APC can still react to upstream WNT cues, and that receptor-based WNT signals differ from downstream activation^32,33^.

Together, our findings support a conceptual model in which oncogenic context dictates how WNT signaling autonomy is achieved in colorectal cancer. In BRAF/RNF43-mutant tumors, receptor sensitization creates a state of heightened ligand dependence that can be resolved by activation of epithelial Wnt7b secretion, establishing a local autocrine WNT signaling loop. This self-supplied ligand source sustains β-catenin–dependent signaling and tumor cell survival in the absence of exogenous WNTs. In contrast, APC-mutant tumors do not require WNT ligands for proliferation, yet ligand availability still modulates transcriptional programs, indicating that extracellular WNT signals continue to tune tumor cell state independently of growth control.

Despite previous evidence of epithelial WNT secretion in CRC^14^, our study provides the first mechanistic dissection of an autocrine WNT signaling loop in CRC and identifies *Wnt7b* re-expression as a key event. In the CRC context, Wnt7b signals via canonical, β-catenin dependent WNT activation, although it has also been implicated in β-catenin independent WNT activation in pancreatic and prostate cancer^69,76^. Further, we defined the spatial dynamics of Wnt7b secretion. Short-range paracrine secretion proved insufficient to sustain proliferation of neighboring cells in a chimeric organoid assay. This is supported by prior work demonstrating that WNT ligands do not diffuse freely, remain largely membrane-associated and that autocrine/juxtacrine signaling dominates in epithelial niches^6^.

Epithelial re-expression of *Wnt7b* and other WNT ligands is not unique to our *in-vitro* model but has also been described *in-vivo*. In line with our patient data, *WNT7B* was shown to be upregulated in human CRC and is associated with increased metastasis^71,77^. Apart from CRC, WNT7B has also been implicated in gastric^78^, breast^79^, prostate^80^, lung^81^ and pancreatic cancer^67,69,82^. Especially in pancreatic ductal adenocarcinoma (PDAC) autocrine WNT7B secretion is an established mechanism enabling WNT self-sufficiency^82,83^.

On a regulatory level, we reveal that early *Wnt7b* expression in Apc and Rnf43/Znrf3 mutated organoids coincides with increased chromatin accessibility at the *Wnt7b* promoter which is bound by the transcription factor Rfx7. These findings nominate Rfx7 as a candidate regulator of *Wnt7b* activation during early oncogenic transformation. This regulatory logic contrasts with a recent study in gastric cancer implicating Smad2/3-dependent control of Wnt7b downstream of oncogenic KRAS^84^. In our system, *Wnt7b* induction occurs independently of KRAS mutations and persists following *Tgfbr2* loss, indicating that intact TGFβ signaling is not required. Together, these observations argue against a primary regulatory role for Smad2/3 signaling and suggest that distinct, context-specific transcriptional programs drive Wnt7b activation in colorectal cancer.

It is likely that the concomitant expression in both Apc and Rnf43/Znrf3 mutated organoids reflects a shared regulatory program upstream of *Wnt7b*. Notably, *Wnt7b* is upregulated in both high- and low-WNT contexts and its expression is not influenced by WNT and R-spondin supplementation. Thus, it is unlikely that ligand availability or WNT activity itself directly control its expression. Further, the lack of activating mutations in MAPK and TGFβ pathways in RZ and A organoids suggests that the Wnt7b regulatory programs differ from other oncogenic programs frequently active in CRC.

Wnt7b is expressed during development and regeneration of multiple endoderm- and ectoderm-derived organs^85–90^. Since oncoembryonic programs are often hijacked by cancer cells to acquire plasticity^91^, such programs are likely candidates controlling Wnt7b. In this framework, epithelial Wnt7b induction may reflect activation of a broader developmental program, with functional consequences that differ depending on the underlying genetic context, being essential for growth in BRAF-mutant CRC while largely dispensable in APC-mutant tumors.

Although the relative contribution of stromal versus epithelial WNT sources remains unresolved, Porcn inhibition effectively suppresses WNT-dependent tumors by blocking both epithelial and stromal compartments, as shown in multiple cancer models^56,92,93^. Porcn inhibitors are currently in clinical trials for RSPO-fusion positive and RNF43-mutant tumors and show promising preclinical efficacy, but their clinical use is limited by on-target toxicity in other tissues as they inhibit WNT secretion systemically^94–97^. Our finding that Wnt7b is the only WNT expressed in our BRAF-mutant model is therefore highly clinically relevant. This is corroborated by a recent CRISPR screen that discovered *WNT7B* as a critical vulnerability in a MSI PDO with striking resemblance of to our model (*BRAF*, *RNF43*, *TGFBR2*, *TP53* mutations)^71^. Targeting WNT7B, which is not expressed in the healthy colonic tissue, could offer a more specific therapeutic window than blocking the secretion of all WNT ligands systemically. Coupled with targeted delivery strategies, this specific inhibition of WNT7B could be a promising therapeutic approach.

In summary, we demonstrate that epithelial plasticity in ligand secretion – rather than just receptor rewiring or downstream activation – can create self-sustaining autocrine circuits that redefine classical notions of pathway dependence. Such adaptations may represent a general adaptive strategy across cancer types to achieve signaling autonomy. By uncovering a *Wnt7b*-driven autocrine loop as a key determinant of Braf-mutant CRC growth, we redefine how WNT plasticity fuels malignant progression and identify a tractable entry point for therapeutic intervention.

### Limitation of the study

A limitation of our study is the absence of stromal and immune compartments in the organoid systems used. While our data demonstrate that epithelial autocrine WNT secretion is sufficient to sustain growth *in vitro*, *in vivo* studies have shown that stromal WNT ligands, including CAF-derived Wnt2 and Wnt5a, also contribute to CRC progression^98,99^. Whether paracrine stromal WNT signals can compensate for, or override, epithelial autocrine WNT secretion, particularly in ligand-dependent tumor contexts, remains unresolved. Prior studies addressing tumor-intrinsic WNT secretion in APC-mutant models have yielded divergent results ^100,101^, underscoring strong context dependence. Future studies using genetically defined multicellular systems and *in vivo* models will be required to delineate how epithelial and stromal WNT sources are integrated during CRC progression.

## Materials and methods

### Mice

To enable conditional *Wls* recombination, R26-mTmG mice^102^ were crossed with *Villin*-Cre^ERT2^; *Wls^fl/fl^* mice^10^ to generate the R26-mTmG; *Villin*-Cre^ERT2^; *Wls^fl/fl^* strain. Organoids were isolated from a 10-week-old female R26-mTmG; *Villin*-Cre^ERT2^; *Wls^fl/fl^* animal.

Animal experiments were performed in accordance with Swiss Guidelines and approved by the Cantonal Veterinary Office Zürich, Switzerland. Mice were housed under specific-pathogen free conditions in a 12 h-12 h light-dark cycle. Chow and water were provided ad libitum.

### Culturing of murine organoids

R26-mTmG; *Villin-*Cre^ERT2^*; Wls^fl/fl^* wildtype colonic organoids were extracted and cultured in 40 μl Matrigel domes (Corning) as described previously^103^. For WT organoids, Basal medium consisting of Advanced DMEM/F12 (Gibco) supplemented with 10 mM HEPES (Gibco), 1X GlutaMax (Gibco), 1% penicillin–streptomycin (Gibco), 1X B27 supplement (Gibco), 1X N2 supplement (Gibco), 100 µg/mL Primocin (InvivoGen), 1 mM N-acetylcysteine (Sigma-Aldrich). In addition, Basal medium was supplemented with the following murine niche factors: 0.5 nM WNT surrogate (ImmunoPrecise), 50 ng/mL EGF (Life Technology), 500 ng/mL R-spondin 1 (Peprotech) and 100 ng/mL Noggin (Sigma). The combinations of niche factors used for the different versions of cancer organoids are depicted in the table below. Depending on organoid type, organoids were passaged every 2-4 days by mechanical dissociation. Brightfield images and fluorescent images were taken on a Revolve microscope (Echo).

| Organoid type | Niche factors used |
| --- | --- |
| WT | WNT surrogate, R-spondin 1, EGF, Noggin. For WNT starvation only EGF and Noggin were used. |
| A, RZ | EGF, Noggin |
| BRZ, BMRZ, AK, AKP | Noggin |
| BRZPS, BMRZT, BMRZTP, AKPS | - |
| BMRZT *Wls*<sup>-/-</sup>, BMRZTP *Wls*<sup>-/-</sup>, BMRZTP *Wnt7b*KO | WNT surrogate, R-spondin 1 |

### EdU proliferation assay

To assess cell proliferation, organoids were processed using the Click-iT Plus EdU Alexa Fluor 647 Flow Cytometry Assay Kit (Thermofisher). Organoids were labelled by adding EdU (final conc. 10 μM), followed by 1.5 h incubation at 37°C to allow for EdU incorporation. Following incubation, organoids were harvested and dissociated into single cells using TrypLE express enzyme (Gibco). Subsequently, EdU labeling was performed according to the manufacturer’s recommendations. Finally, cells were stained with DAPI (Thermofisher, 1:1000) and analyzed immediately on a FACSymphony 5L (BD) analyzer. Data was analyzed in FlowJo (BD Biosciences).

### DNA/RNA extraction

Organoid DNA from at least two filled 40 μL domes was extracted using the NucleoSpin Tissue kit (Macherey-Nagel) according to the manufacturer’s recommendations. Organoid RNA from at least 4 filled 40 μL domes was extracted using the NucleoSpin RNA kit (Macherey-Nagel) according to the manufacturer’s recommendations, including the optional DNase digest. Both RNA and DNA concentrations were measured using the NanoDrop 1000 (Witec).

### Tamoxifen-induced recombination of *Wls^fl/fl^* organoids

After passaging, R26-mTmG*; Villin-*Cre^ERT2^*;Wls^fl/fl^* BMRZT, BMRZTP, and AKPS organoids were treated with 1 μM 4-OHT for 16 h to induce Cre^ERT2^-mediated recombination. Wells were then washed with prewarmed PBS, replenished with fresh medium, and cells were allowed to expand. Recombination at the R26-mTmG locus was used as a proxy for *Wls* deficiency and GFP⁺ cells were isolated by FACS (FACSAria III 5L sorter, BD) at the flow cytometry facility of the University of Zurich. Cells were subsequently maintained in medium supplemented with WNT surrogate and R-spondin 1.

### MSI testing

To test microsatellite instability (MSI), length of five previously published microsatellite loci^51,52^ was assessed. DNA from WT and BMRZTP organoids was extracted as described above and amplified using labelled primers with the Q5 High fidelity DNA polymerase (New England Biolabs) according to the manufacturer’s recommendation (Primers are indicated in the table below). Labelled amplicons where then sent for fragment length analysis by capillary electrophoresis at Microsynth (Balgach, Switzerland) using a DS-30_D Filterset and GeneScan 400 HD ROX size standard. Data was analyzed using GeneMapper software (Applied Biosystems).

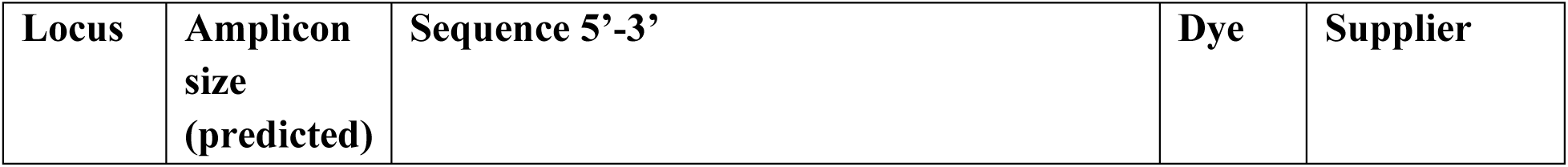
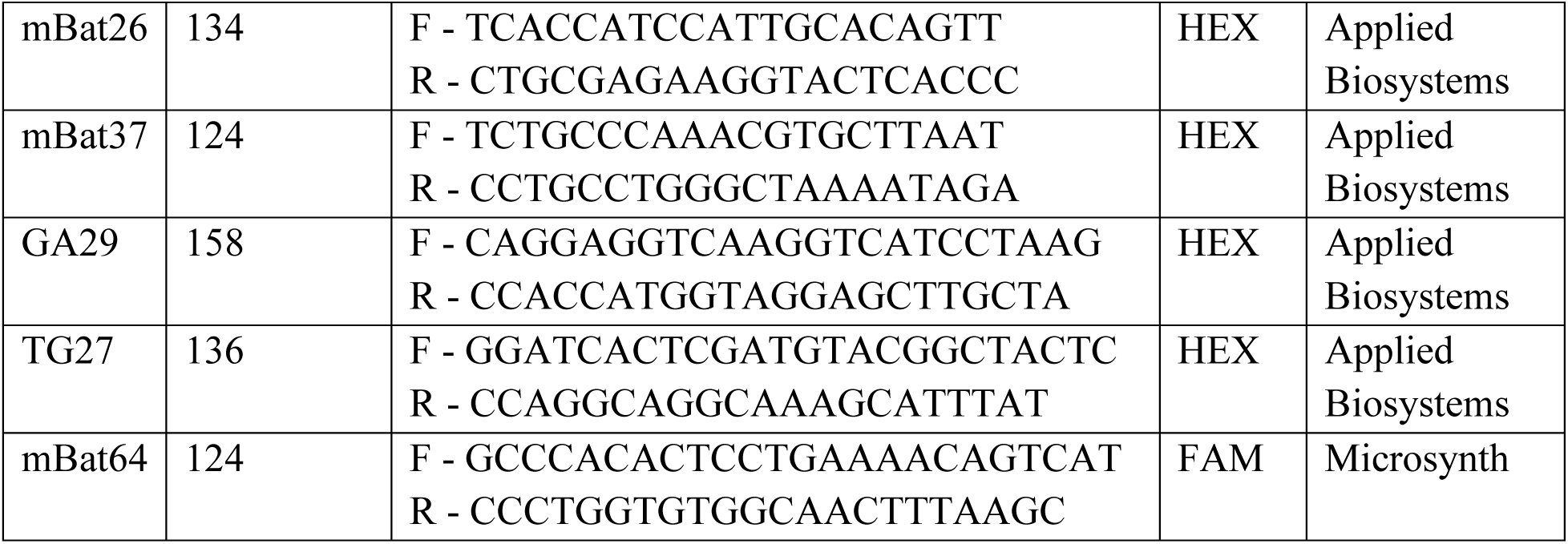

### Reverse transcription and RT-qPCR

To generate cDNA, 500 ng of RNA was reverse-transcribed using the cDNA synthesis kit (Takara Bio) according to the manufacturer’s recommendations. The primers used for RT-qPCR are listed below.

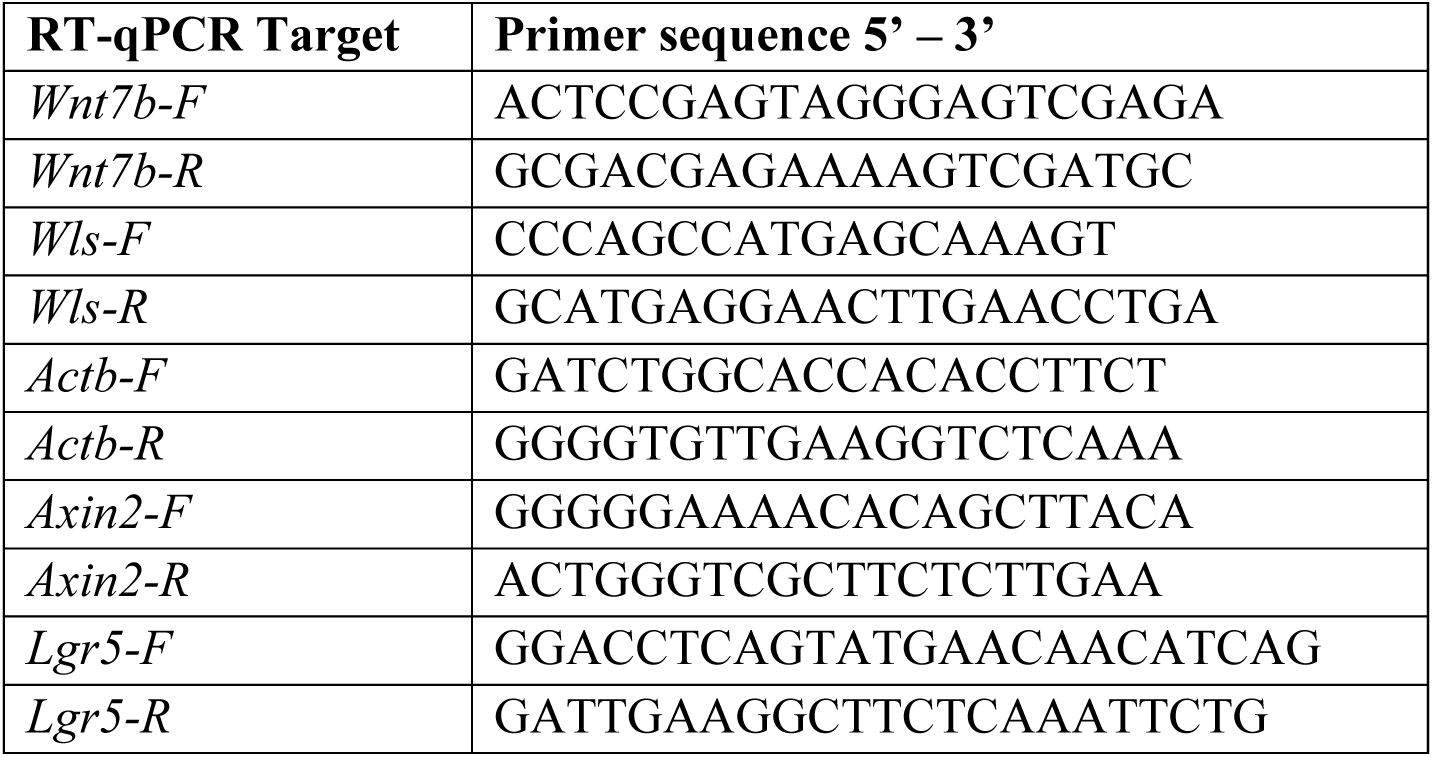

Gene expression was determined by RT-qPCR using the PowerTrack SYBR Green Master Mix (Thermofisher) on the QuantStudio3 system (Applied Biosystems). The samples were analyzed in technical triplicates and the average cycle threshold values were normalized to *Actb* expression using the ΔΔCT method^104^.

### Cloning and plasmids

To generate the mutant *Braf^V637E^* overexpression construct, the murine *Braf* coding sequence was amplified from cDNA extracted from wildtype organoids. The V637E substitution, corresponding to human *BRAF^V600E^*, was introduced by site-directed mutagenesis. The resulting amplicon was subcloned into a lentiviral bicistronic expression construct by Gibson assembly (PGK-Blasticidin-T2A-*Braf^V637E^*). Phenotypic similarity of human and murine isoforms was assessed by pERK western blot in HEK cells transfected with the respective constructs.

The plasmid pEF1a-mMLH1dn (a gift from Dr. David Liu; Addgene plasmid #174825) was modified by insertion of a puromycin resistance gene under the control of a PGK promoter via restriction site cloning. The pLenti-PGK-KRAS(G12V) (Addgene # 35634) construct was a kind gift from Dr. Daniel Haber.

### Lentiviral production and spinoculation

To produce lentivirus, HEK cells were seeded in lentiviral packaging medium (5% FBS + 0.2 mM Sodium Pyruvate in OptiMEM). Transfection was performed using Lipofectamine 3000 (Thermofisher) according to the manufacturer’s recommendation. In combination with the lipofectamine reagents, 9 μg of lentiviral expression vector, 2.25 μg pRev (Addgene 12253), 2.25 μg pVSV (Addgene 12259) and 6.75 μg pMDL (Addgene 12259) were used. Lentiviral supernatant was harvested after 16, 40, and 64 hours, with fresh packaging medium added after each collection. Following the last collection, lentiviral supernatant was filtered through a 0.45 μm filter (Cytiva), and concentrated using an Amicon Ultra-15 Centrifugal Filter unit (Merck Milipore) to 1 mL and stored in 100 μL aliquots at −80°C.

For lentiviral transduction, organoids were harvested, washed once and dissociated to single cells with TrypLE at 37°C for 7 min. The reaction was quenched with DMEM + 10% FBS (Gibco) and cells were centrifuged for 5 min at 300 x g at 4°C. Supernatant was aspirated and cells were resuspended in 50 μL lentiviral solution + 450 μL of Basal medium supplemented with the required niche factors, 10 μM ROCK inhibitor, 5 μM nicotinamide and 4 ng/μL Polybrene. Cells were subsequently spinoculated for 1 h at 600 × g in a prewarmed centrifuge (37 °C), then returned to the incubator (37 °C) for recovery for 2 h. Finally, cells were pelleted, resuspended in Matrigel and plated in 40 μL domes. After outgrowth, cells were selected for at least 2 passages using the respective antibiotics.

### RNP-mediated CRISPR-Cas9 knockout

To knock out *Apc*, *Tp53*, *Smad4*, *Rnf43*, *Znrf3* and *Tgfbr2* sgRNAs were ordered from Integrated DNA technologies (IDT). Sequences are indicated in the table below. The sgRNAs were a combination of both tracrRNA and crRNA (Alt-R system, IDT). All organoids in this study were generated starting from WT colonic organoids extracted from R26-mTmG; *Villin-*Cre^ERT2^*; Wls^fl/fl^* animals. To prepare RNP complexes 1.34 μL of sgRNA (100 μM) and 1.67 μL of S.p. Cas9 Nuclease V3 (IDT) were mixed and incubated for 10 min at RT. After incubation, 5.33 μL of OptiMEM (Gibco) were added to the RNP solution. In the meantime, organoids growing in 2-3 full domes of 40 μL Matrigel (Corning) were harvested, washed and dissociated into a single cell solution by incubation in 1 mL 1X TrypLE (Gibco) at 37°C for 7 min. After inactivation with DMEM (Gibco) supplemented with 10% FBS (Gibco), cells were washed twice with OptiMEM (Gibco) and centrifuged for 5 min at 290 x g at 4°C. Supernatant was carefully aspirated and cells were resuspended in 94 μL of OptiMEM (Gibco) supplemented with 5 nM nicotinamide (Sigma) and 10 μM ROCK inhibitor (Y-27632, Stem Cell Technologies).

The cell suspension was then mixed with the 8.33 μL RNP solution prepared previously and 1 μL of Electroporation enhancer (100 μM, IDT). Prior to electroporation, exactly 100 μL of the finals cell suspension was added into electroporation cuvettes (EC-002S, Nepagene) and cells were electroporated in a NEPA21 electroporator (Nepagene) using the following settings: Poring pulse voltage =175V, Length = 5 ms, Interval = 5 ms, Number of pulses = 2, Decay rate = 10%, Polarity = +; Transfer Pulse voltage = 20V, length = 50 ms, Interval = 50 ms, Number of pulses = 5, Decay rate = 40%, Polarity = +/−. Impedance of the suspension was expected to be between 30 – 55 Ohm. After electroporation, 0.5 mL of prewarmed Advanced DMEM/F12 (Life Technologies) including the respective niche factors for the different organoid types + 5 nM nicotinamide (Sigma) and 10 μM ROCK inhibitor (Y-27632, Stem Cell Technologies) was added and the solution was pipetted up and down to break formed foam. The cell suspension was then transferred to a 48-well plate (Sigma) and placed into the incubator at 37°C for 2 h. Finally, cells were washed once and plated using double the amount of Matrigel from which they were harvested.

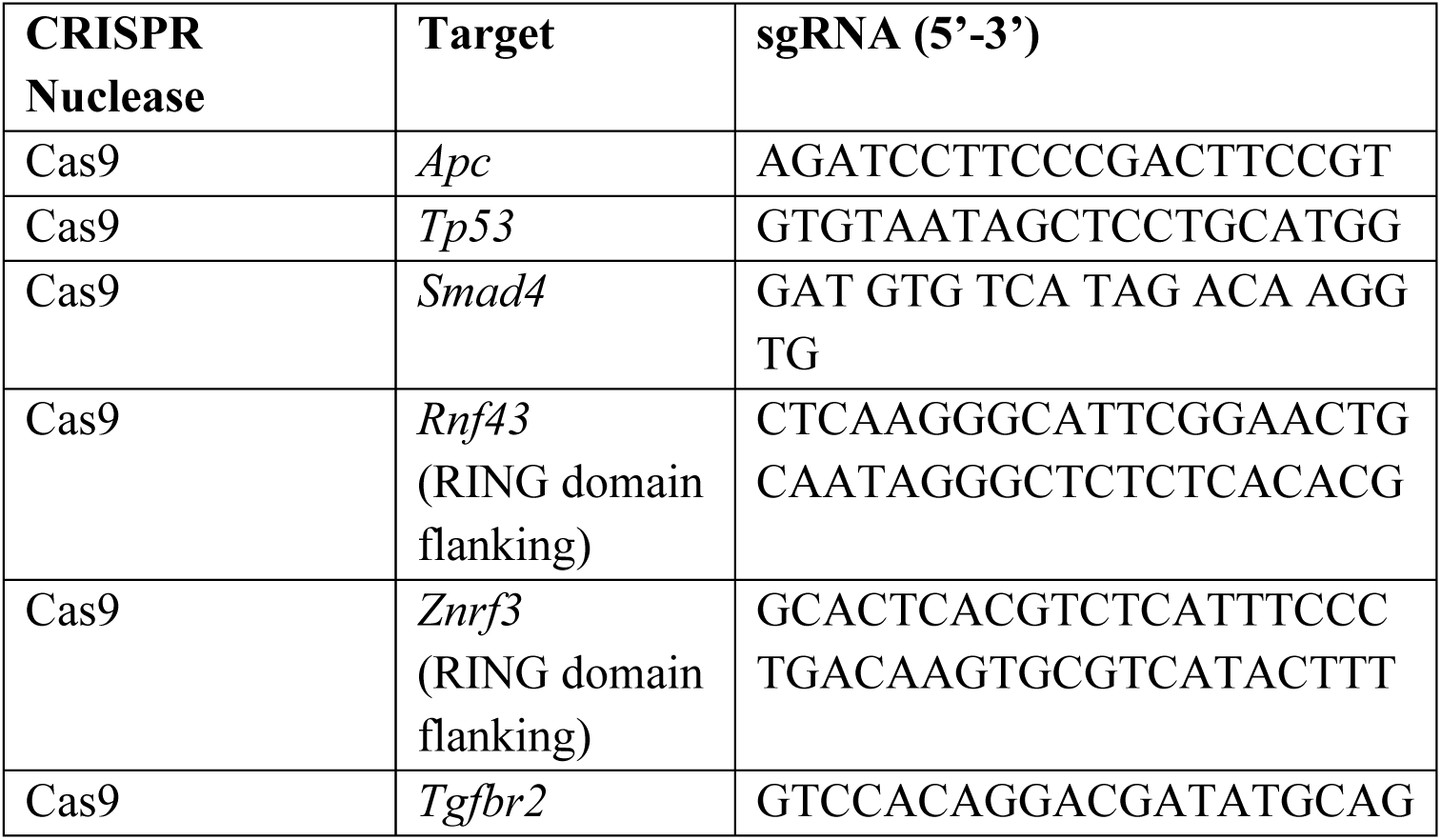

Following metabolic selection, the targeted loci were amplified by PCR. For *Rnf43* and *Znrf3,* successful deletion of the RING-Domain was assessed by Gel electrophoresis. For the other targets, PCR products were cloned into a pGEM vector with the pGEM-T Easy Vector system (Promega) and at least 28 colonies per gene were sequenced by E.coli NightSeq (Microsynth). To assess mutational status, sequences were aligned in CLC Main workbench 7 (Qiagen).

### Genotyping primers CRISPR-Cas9

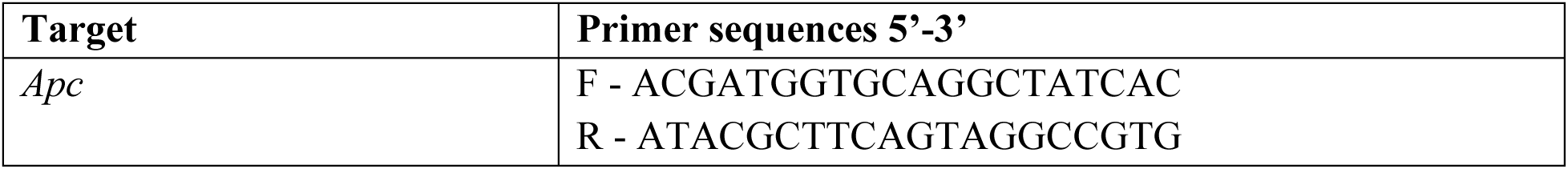
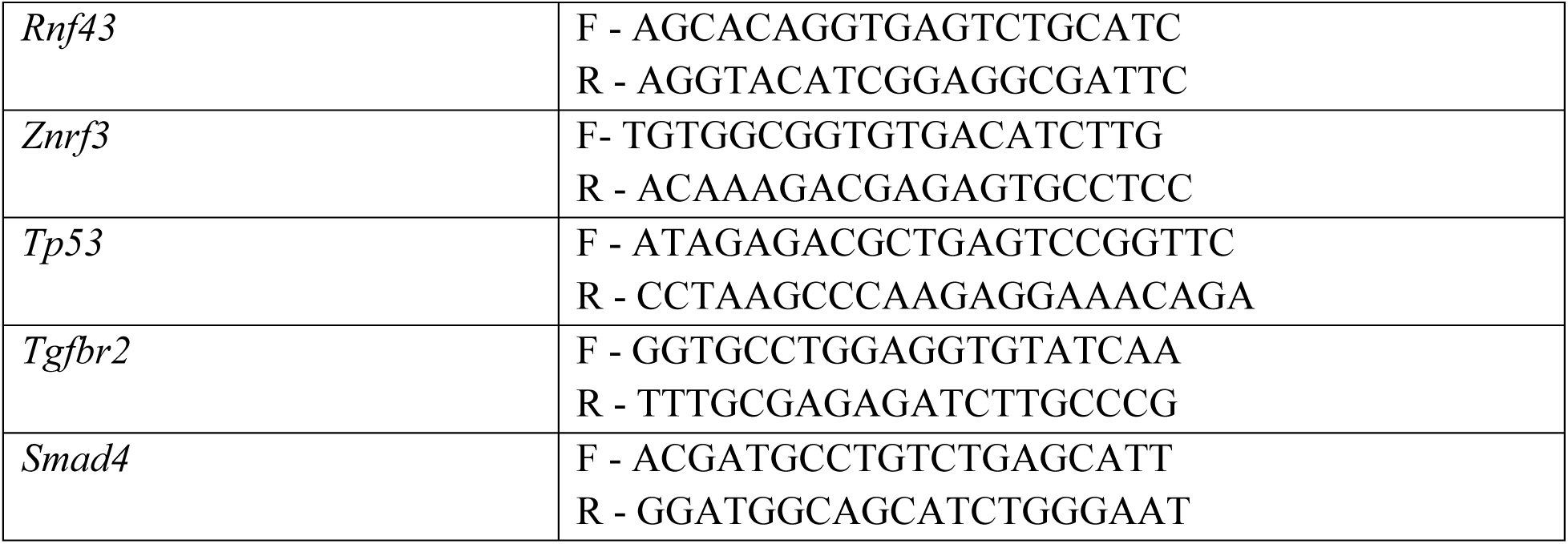

### EnAsCas12a knockout of *Wnt7b* in BMRZTP organoids

BMRZTP *Wnt7b* knockout organoids were generated using enAsCas12a^105,106^ in combination with Pol II delivered crRNA-arrays^107^. The necessary spacer sequences, targeting *Wnt7b* and intergenic regions, were obtained using the guide design tool CRISPick^106,108^ with the mouse GRCm38 reference genome and enAsCas12a CRISPRko mode. SgRNAs are depicted in the table below. For *Wnt7b* three spacers were chosen (efficacy scores > 0.8 without off-targets) to construct 3-mer pre-crRNA arrays for single knockouts. Similarly, 6-mer pre-crRNA arrays were designed for non-targeting and intergenic regions. The control array was cloned into a 2^nd^ generation lentiviral vector, which expressed the array under the EF1a promoter together with GFP and Puromycin (EF1a-EGFP-2A-Puro-Triplex-pre-crRNA array-WPRE)^107^. In parallel, the *Wnt7b* array was cloned into a 2^nd^ generation lentiviral vector, which expressed the array under the EF1a promoter together with mRuby and Zeocin (EF1a-mRuby3-P2A-Zeocin-Triplex-pre-crRNA array-WPRE)^107^. Both constructs were compatible with enAsCas12a-2A-Hygro (pRDA_174, Blasticidin replaced with Hygromycin, Addgene, #1346476^106^) in BMRZTP organoids.

### EnCas12a sgRNAs

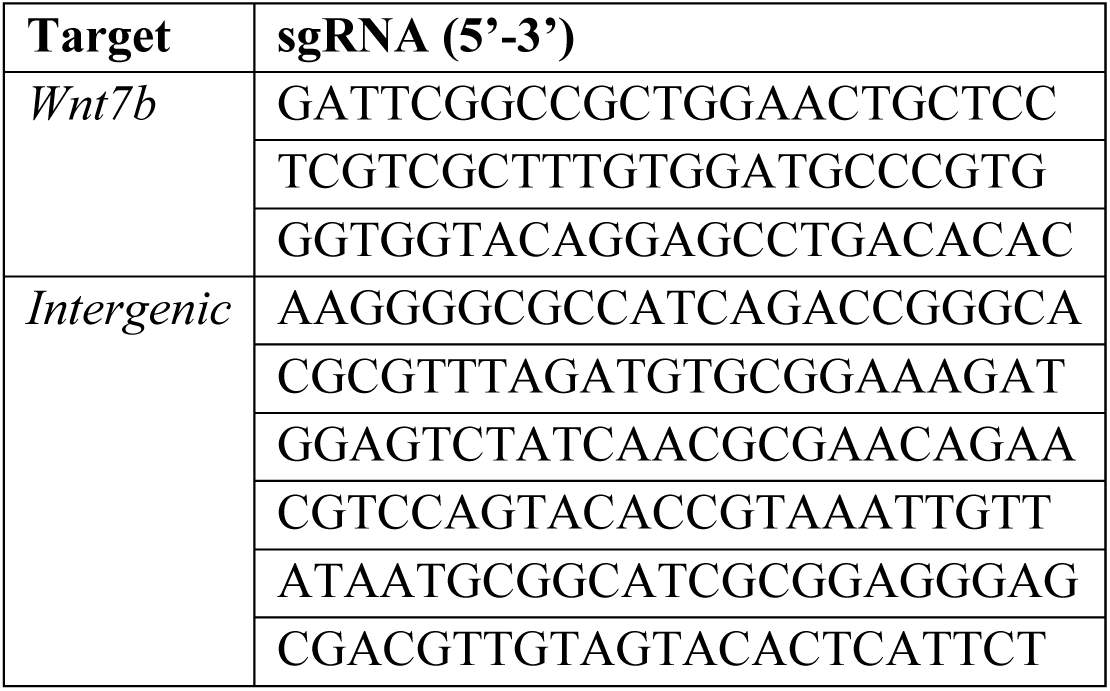

Lentiviral transduction was performed as described above and transduced organoids were selected at least for two passages with 400 μg/mL Zeocin (Invivogen) and 200 μM Hygromycin B Gold (Invivogen). In the case of CTRL organoids, EGFP^+^ organoids were sorted by FACS (FACSAria III 5L sorter, BD) at the flow cytometry facility of the University of Zürich. To confirm genomic editing of *Wnt7b* in BMRZTP organoids, regions around the 3 targeted loci were amplified by PCR (Primer table below) and products were cloned into a pGEM vector with the pGEM-T Easy Vector system (Promega). At least 24 colonies per locus were sequenced by E.coli NightSeq (Microsynth). To assess mutational status, sequences were aligned in CLC Main workbench 7 (Qiagen).

### Genotyping primers enAsCas12a

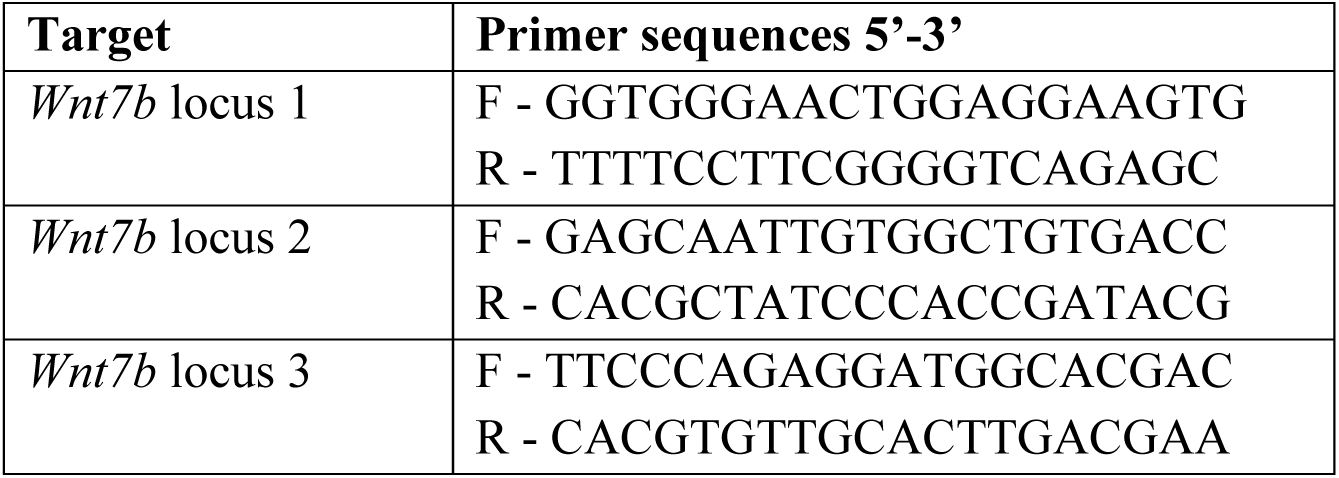

### Metabolic selection and inhibitor treatments

Following RNP electroporation or lentiviral transduction, cells harboring the desired mutations/ expressing the desired transgenes were selected as follows: For the selection of *Apc*-mutated cells, organoids were cultured in medium lacking WNT-surrogate and R-spondin 1. To select for cells harboring *Rnf43*/*Znrf3* mutations, cells were cultured without R-spondin 1 or without both WNT surrogate and R-spondin 1. To select cells transduced with the *Bra^fV635E^* construct, organoids were cultured in medium lacking EGF and additionally supplemented with 1μM the EGFR inhibitor (HDS029, Santa Cruz) and 5ng/μl Blasticidin (Sigma) for at least 3 passages. Cells transduced with *KRAS^G12V^*were selected by removal of EGF from the culture medium and supplementation with 100 μM Hygromycin (Sigma) for at least 2 passages. *Mlh1*dn expressing cells were selected with 1.5 μg/mL Puromycin (Sigma) for at least 2 passages. *Smad4* and *Tgfbr2* mutant cells were selected by withdrawing Noggin and the addition of 25 ng/μL TGFβ (Peprotech). *Tp53*-mutant cells were selected by 10 μM Nutlin-3 (Cayman chemicals) for at least 4 passages. For Porcupine inhibition experiments, 1 μM IWP-2 (Stemcell) was used for 2 passages.

### Longitudinal FACS analysis of *Wls*^+/+^ and *Wls*^−/−^cells

BMRZT organoids cultured in +WR were split into six wells of a 6-well plate (Sigma-Aldrich) with 7 domes per well. 24 h after passaging, BMRZT organoids were treated with 1μM 4-OHT for 24 hours to induce partial *Wls* recombination. The resulting chimeric organoids consisted of *Wls*^+/+^, TdTomato^+^ cells and *Wls*^−/−^, GFP^+^ cells. Following the 4-OHT treatment, the wells were washed once with PBS (Gibco) and +WR medium was added to three wells. To the other three wells, fresh −WR medium was added. 24 hours after the medium change, two out of three wells per condition (+WR or −WR) were harvested and dissociated to a single cell suspension. Cells were stained with DAPI (Thermofisher, 1:1000) and subsequently the percentage of live TdTomato^+^ and GFP^+^ cells was determined by flow cytometry using a FACSymphony 5L (BD) analyzer. This first analysis corresponded to timepoint Day 1. The same day, the remaining organoids were split and kept in either +WR or – WR medium. This procedure was repeated in identical manner on Day 15 and Day 22 of the experiment.

### Luciferase assay

The proximal murine *Wnt7b* promoter region (3kb) was cloned into a pGL3 basic vector (Addgene #212936), directly upstream of Firefly luciferase, to generate a reporter construct. As control for normalization, the pRL Renilla Luciferase vector (Promega, E2231) was used. A self-cloned CMV-EGFP-H2B vector was used as lipofection control. For overexpression of candidate transcription factors, the constructs listed in the table below were used.

To measure luciferase activity, 10’000 HEK293T cells were seeded in DMEM medium + 10% FBS (Gibco) per one well of a 96-well plate. Per condition, three technical replicates (wells) were prepared. After 24 hours, cells were transfected with 17 ng of reporter vector (pGL3), 3.4 ng Renilla vector (pRL) and 34 ng of transcription factor construct per well using Lipofectamine 3000 reagent according to the manufacturer’s recommendation. 24 hours after lipofection, medium was removed and cells were washed once with PBS (Gibco). Cells were then lysed using 1 x passive lysis buffer diluted in water (Promega, E194A) and incubated at −80°C for 30 min. Subsequently, 10 μL of the lysed cell suspension was transferred to a fresh, white 96-well plate. Luminescence of Firefly and Renilla luciferase were then measured successively after adding 50 μL of luciferase assay reagent II (Promega, E195A) and Stop&Glo reagent (Promega, E641A) in a GloMax-Multi microplate reader (Promega).

### Plasmids encoding candidate transcription factors

| Transcription factor | Addgene number |
| --- | --- |
| Rfx7 | #142554 |
| Zfx | #143026 |
| Nr2c1 | #142285 |
| Hif1a | #142638 |
| Arnt | #143322 |
| Runx2 | #143554 |

### Tissue fixation, paraffin embedding and generation of paraffin sections

Human colorectal cancer samples were fixed for 24 hours at room temperature in 4% paraformaldehyde in PBS (Gibco). Tissues were then dehydrated through graded EtOH. Subsequently, tissues were cleared with Xylene. Samples were then infiltrated with molten paraffin and embedded in paraffin blocks. After solidification, blocks were sectioned using a HM355s microtome (Thermofisher).

### Immunofluorescent staining

Paraffin sections were deparaffinized and rehydrated by serial washes through Xylene and graded EtOH to PBS. Next, sections were submerged in citrate buffer (pH = 6.0) and antigen retrieval was performed using a high-pressure histological microwave (HistosPro, Milestone) at 110°C. After retrieval, samples were cooled down and washed once in PBS before proceeding with antibody staining.

Tissue was blocked with 5% BSA (Roche) + 0.2% Tween 20 (Sigma) in PBS (Gibco) followed by 1 h incubation at room temperature with primary antibodies. The following primary antibodies were used: anti-Wnt7b (Invitrogen, Cat. No. PA-103480, 1:200), anti-E-cadherin (Biotechne, AF748, 1:200) diluted in 3% BSA. Consequently, samples were washed three times with PBS (Gibco) and incubated with the secondary antibodies Donkey anti-Rabbit IgG Alexa Fluor 488 (Invitrogen A21206, 1:500), Donkey anti-goat IgG Alexa Fluor 647 (Invitrogen, A32795, 1:500) and DAPI (Thermofisher) diluted in 3% BSA for one hour at RT in the dark. After washing with PBS (Gibco), sections were mounted with Dako fluorescence Mounting Medium (Agilent) and left to dry overnight in the dark. For imaging, a Leica SP8 inverse confocal microscope was used. Images were processed in FIJI (ImageJ 1.53c).

### *Wnt7b*KO rescue experiment and quantification of organoid diameters

To rescue the phenotype of *Wnt7b*KO organoids, following passage, organoids were supplemented with WNT surrogate (0.5 nM) and R-spondin1 (500 ng/ml) (+WR), WNT surrogate (0.5 nM) (+W), medium lacking WNT surrogate and R-spondin (−WR), R-spondin (500 ng/ml) (+R), CHIR99021 (10 μM) (StemCells, Cat. No. 72054), Wnt3a (400 ng/ml) (Biotechne, Cat. No. 1324-WN-010) supplemented with R-spondin1 or Wnt5a (250 ng/ml) (Biotechne, Cat. No. 645-WN-010) supplemented with R-spondin1 (500 ng/ml). Organoids were cultured in 50 μL domes and medium was replaced daily. After 48 hours, organoids were passaged a second time. 48 hours after the second passage, multiple images per dome were acquired using a Revolve microscope (ECHO). This experiment was repeated three times to generate three biological replicates. Subsequently, organoid diameters were measured manually using ImageJ. At least 25 organoids per replicate and condition were analyzed.

### Immunoblotting

For immunoblot analysis cell were harvested and lysed in 150 μL cold RIPA buffer supplemented with 1 X complete protease inhibitor cocktail (Roche) and phosphatase inhibitor (Roche). After 15 min incubation on ice, samples were centrifuged at 15’000 x g for 10 min on 4°C. The supernatant was transferred to a new tube and protein concentrations were determined using a Micro BCA Protein assay kit (Thermofisher). Samples were separated by SDS-PAGE using 4-12% Bis-Tris Gels (Invitrogen). Due to similar molecular mass of pERK and Tubulin, two separate gels were prepared. Proteins were then blotted onto a 0.45μm Polyvinylidene difluoride (PVDF) membrane (Amersham Hybond) and incubated with the primary antibodies anti-alpha tubulin (Sigma, T6199, 1:2000) and anti-p44/42 (Cell signaling, D13.14.4E, 1:2000) overnight at 4 °C. Subsequently, the following HRP-conjugated secondary antibodies were used: Goat anti-rabbit IgG (Jackson, Cat. No. 111-035-144, 1:5000) and Goat anti-mouse (Invitrogen, G-21040, 1:5000). After washing, blots were incubated in a 1:1 mix of WesternBright (Advansta) for 5 min and subjected to imaging using a FUSION FX6 gel imager (Vilber).

### Bulk RNA sequencing

Before RNA collection, all sequenced organoids were cultured in −WREN medium for 48h with the exception of Wt_comp that were cultured in complete medium and A and RZ organoids which were cultured + EGF and + Noggin before RNA collection. Wt_wren organoids were cultured 24 h + WREN followed by 48 h −WREN. Organoids strictly dependent on exogenous WNTs (BMRZT_Wls, BMRZTP_Wls) were split and cultured for 48 h in −WREN medium before RNA collection. Likewise, BMRZT_Porcni organoids were cultured 48 h in −WREN supplemented with 1 μM IWP-2 before harvesting. RNA from colonic organoids was extracted using the NucleoSpin RNA Kit (Machery-Nagel) according to the manufacturer’s recommendations including DNase digestion. RNA concentration was measured using the NanoDrop (Thermofisher). Sequencing was performed by the Functional Genomics Center Zurich. Extracted RNA was prepared for sequencing using the Illumina TruSeq Stranded mRNA Library Prep assay following the manufacturer’s protocol. Sequencing was performed on the Illumina NovaSeq X with a paired end 150bp read configuration.

RNA sequencing data was analysed using the SUSHI framework^109^, which encompassed the following steps: Inspection of read quality using FastQC, and removal of sequencing adaptors using Fastp^110^; alignment of the RNA-Seq reads utilizing the STAR aligner^111^ with the Ensembl murine genome build GRCm39 as reference^112^; counting of gene-level expression values using the featureCounts function of the R package Rsubread^113^; Assessment of differential expression using the generalized linear model as implemented by the DESeq2 Bioconductor R package^114^ and Gene Ontology (GO) term pathway analysis using the hypergeometric over-representation test via the enrichGO function of the clusterProfiler Bioconductor R package^115^. All R functions were executed on R version 4.4.2 (R Core Team, 2020) and Bioconductor version 3.20. Bioinformatic data handling was performed by bioinformatician Dr. Peter Leary at the Functional genomics center Zurich.

For further downstream analysis of overlapping DEGs between multiple conditions, differential expression results from DESeq2 were imported into *SummarizedExperiment* in R Studio. To compute overlaps between conditions with *ComplexHeatmap*^116^ lists of significantly up- and down regulated genes (FDR ≤ 0.05, log2FC thresholds as indicated) were generated.

For custom gene set enrichment analysis (GSEA), gene sets with human symbols were converted to mouse gene symbols using the Mouse Genome Informatics (MGI) homology database (http://www.informatics.jax.org). These custom pathways were tested alongside Gene Ontology biological process terms, with significance assessed using permutation testing and normalized enrichment scores (NES).

### ATAC sequencing

ATAC sequencing was performed as previously described^117^. Briefly, 3 replicates of WT, A and RZ (total sample n = 9) organoids were dissociated into single cells. 50’000 cells per replicate were centrifuged for 5 min at 400 x g and resuspended in ice cold lysis buffer (10 mM Tris-HCl, pH 7.4, 10 mM NaCl, 3 mM MgCl2, 0.1% IGEPAL CA-630) by pipetting up and down. Subsequently, the lysed cell suspension was centrifuged 10 min at 500 x g and the supernatant was decanted. For tagmentation, 25 μL Tagment DNA Buffer, 2.5 μL Tagment DNA Enzyme, and 22.5 μL nuclease-free H2O) (Illumina) were added and the reaction was incubated for 30 min at 37°C. The tagmented DNA was purified with the MinElute PCR Purification Kit (Qiagen) and eluted in 10 μL elution buffer. For preamplification, samples were amplified for 5 cycles with Nextera compatible Multiplex primers (ActiveMotif) and NEBnext high fidelity 2X PCR Master Mix (NEB). The reaction was performed using the following components:10 μL transposed DNA, 10 μL water, 2.5 μL 25 μM Nextera primer 1, 2.5 μL 25 μM Nextera primer 2, 25 μL 2X PCR Master Mix. Consequently, 5 μL of the reaction were used for qPCR to assess the additional number of PCR cycles to avoid oversaturation of the amplification. Between 7 and 11 additional cycles were performed. The final libraries were again purified using the MinElute PCR purification Kit and eluted in 20ul elution buffer before determining the DNA concentration at a Qubit device (Thermofisher). Pooled libraries were sequenced by the Functional Genomics Center Zurich using 2 lanes of the 10B flowcell one the Illumina Novaseq X Plus (150bp paired-end configuration). Sequencing was performed to a target depth of 200 million paired-end reads per sample to ensure sufficient coverage of open chromatin regions across the genome.

### ATAC-seq analysis

Raw sequencing reads were quality controlled and trimmed using fastp v0.23.4^110^. Adapter sequences were removed, and reads were filtered for polyX tails and minimum length requirements (18 nt). Trimmed paired-end reads were aligned to the mouse reference genome (GRCm39, GENCODE Release M37) using Bowtie2^118^. PCR duplicates were marked using Picard MarkDuplicates (http://broadinstitute.github.io/picard), and duplicate-marked BAM files were filtered to generate processed alignments for downstream analysis. Open chromatin regions were identified by calling peaks on individual samples using MACS3^119^. Peak calling was performed independently for each of the 9 samples. Differential chromatin accessibility analysis was performed using the DiffBind package. A consensus peak set was generated across all samples with a minimum overlap requirement of 1 sample. Peaks overlapping ENCODE blacklist regions for mm39 were filtered out. Read counts within peaks were quantified using summarizeOverlaps, and samples were normalized using the RLE (Relative Log Expression) method. Differential accessibility testing was performed using DESeq2 to compare conditions. Peaks with an FDR < 0.05 were considered significantly differentially accessible. Differentially accessible peaks were annotated to genomic features and nearby genes using the ChIPseeker R/Bioconductor package^120^ with the mm39 reference genome annotation (TxDb.Mmusculus.UCSC.mm39.refGene and org.Mm.eg.db). Promoter regions were defined as −1 kb to +1 kb relative to the transcription start site (TSS). Overall chromatin accessibility at TSS was tested using deeptools^121^. Transcription factor binding motif enrichment in differentially accessible regions was analysed using monaLisa^122^ with the JASPAR2020 database. Motif scanning was performed using the matchPWM method from TFBSTools^121^ on peak sequences extracted from the mm39 reference genome (BSgenome.Mmusculus.UCSC.mm39). Both mouse-specific (species 10090) and vertebrate transcription factor motifs from the CORE collection were queried. Enriched motifs were then identified in peaks associated with *Wnt7b*.

### Survival analyses

Kaplan-Meier plots showing overall survival of patients harboring either BRAF or APC mutations were generated using the cBioPortal^123–125^. A time cut-off of 120 months was chosen. Logrank Test was performed on all 1958 patients (BRAF n = 179, APC n = 1779). Kaplan-Meier plots based on WNT7B expression were generated using the “Kaplan Meier Plotter” (https://kmplot.com/analysis/)^226^ on standard settings.

### Statistics

All *p*-values were obtained using Prism (GraphPad Software, v10) software, unless otherwise indicated. In all cases at least 3 biological replicates were used, unless otherwise stated. Information on the statistical tests used can be found in the associated figure legends.

### Use of language tools

Artificial intelligence tools (ChatGPT) were used as a writing tool in the process of crafting this manuscriptIt was employed exclusively for proofreading, language-editing and to improve grammar of the written text. The tool was never used to generate original scientific ideas or to interpret them. All concepts, analyses and conclusions presented in this work were brought forward by the authors, who take full responsibility for the originality of this work.

## Acknowledgments

We would like to thank all former members of the Basler group, especially Konrad Basler and George Hausmann for helpful comments during the preparation of the manuscript. Further we would like to thank Michael Brügger, Erich Brunner, Jamie Little and Giulia Moro for fruitful discussions. Finally, we also would like to thank Glauco Camenisch for his help and advice with the MSI analysis.

## Author contributions

Conceptualization: T.V.

Experimental design: S.R.L, H.F. T.V.

Investigation: S.R.L, T.K, P.L, T.D.

Formal analysis: S.R.L, P.L

Data curation: S.R.L, T.K.

Project administration: H.F., T.V.

Resources: C.P, A.M, R.J.P, R.F, H.F, T.V

Manuscript - initial draft (incl. visualization): S.R.L

Manuscript - editing: S.R.L, H.F., T.V.

## Data availability

RNA sequencing data and ATAC sequencing data generated in this study are deposited in the GEO database

## Supplementary information

**Supplementary Figure 1.**
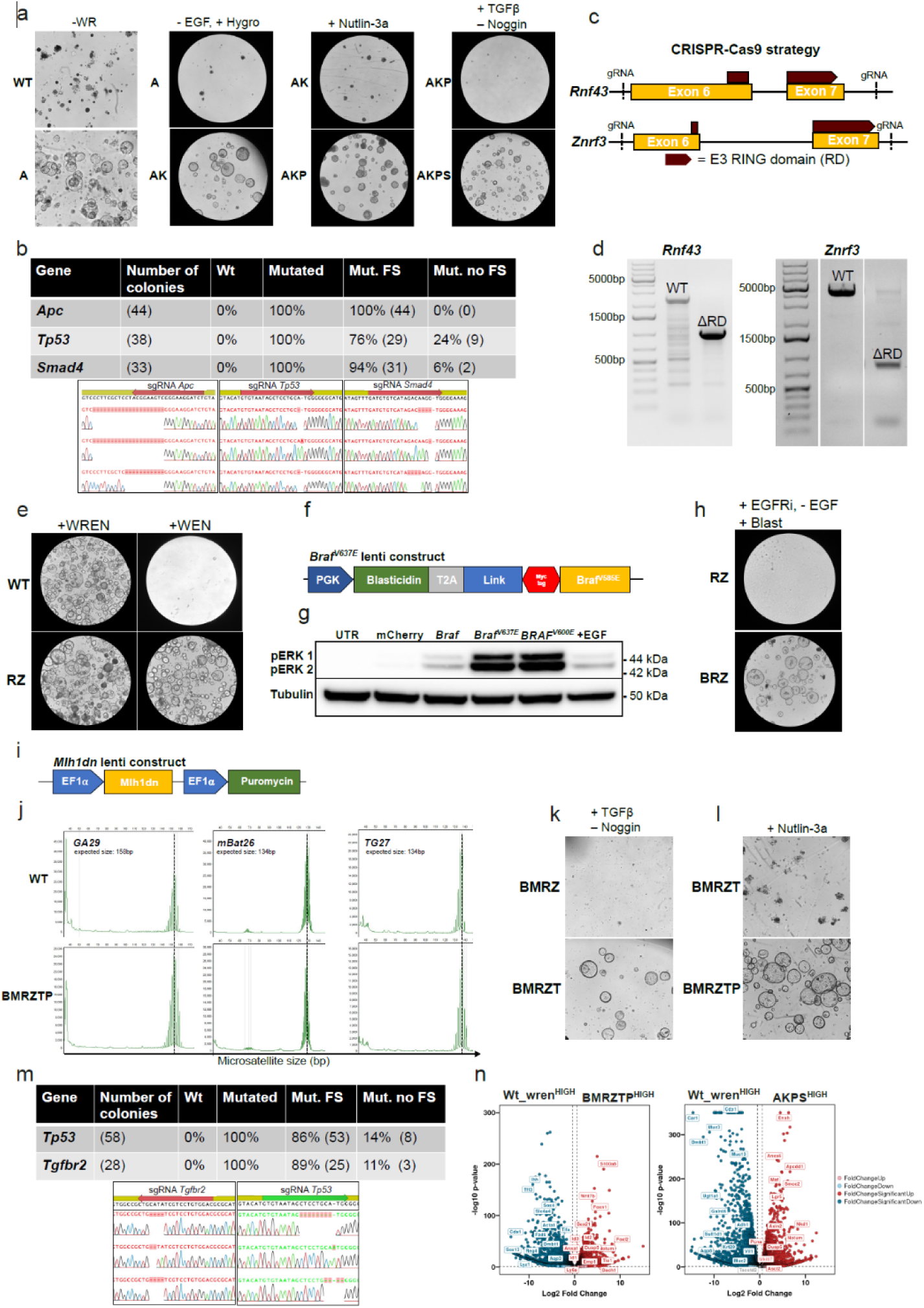
Engineering of AKPS and BMRZTP organoids. a) Metabolic selection of used during the generation of AKPS organoids. From left to right: *Apc* ko. (A) Organoids were selected in medium supplemented with EGF and Noggin by withdrawal of WNT surrogate and Rspo (−WR). Organoids were transduced with a *KRAS*^G12V^ lentiviral construct (AK). Selection was performed by EGF withdrawal and supplementation with Hygromycin. *Tp53* was knocked out in AK organoids (AKP) and mutants were selected with Nutlin-3a. *Smad4* was knocked out in AKP organoids (AKPS) and mutants were selected with TGFβ. In all cases representative images after two passages of selection are shown. b) Sanger sequencing results of the individual mutations in AKPS organoids. Per mutation at least 33 colonies were analyzed. Percentage of mutated colonies with or without frameshift (FS) are shown. Panels below show representative images of the Sanger sequencing results at sgRNA binding sites. c) Schematic visualization of the CRISPR-Cas9 strategy used to remove the RING domain (RD) in *Rnf43* and *Znrf3* (RZ). Per gene, two flanking intronic sgRNAs were used. d) The two panels below show PCR reactions over the targeted region in WT and RZ organoids. Expected fragment sizes of the WT alleles are 2556bp in *Rnf43* and 4063bp in *Znrf3*. Following RD removal, the expected sizes are 1039bp and 772bp for *Rnf43* and *Zrnf3* respectively. e) Growth of WT and RZ organoids 72h after passaging in medium with or without Rspo. f) Schematic visualization of the *Braf*^V637E^-lenti construct. g) PhosphoERK immunoblot performed 48 hours after transfection of HEK293T cells with wildtype or mutant Braf overexpression constructs. All *Braf*-alleles were expressed under a CMV promoter. *Braf*^V637E^ represents the murine isoform of human *BRAF*^V600E^. A plasmid encoding CMV-mCherry was used as negative control. UTR = Untreated transfection control. EGF supplementation served as positive control. h) Selection of RZ organoids transduced with the *Braf*^V637E^ lentiviral construct (BRZ). For selection, the EGFR inhibitor (EGFRi) HDS029 + Blasticidin was used. In addition, EGF was withdrawn from the culture medium. Representative images after two passages of selection are shown. i) Schematic visualization of the dominant negative Mlh1 (Mlh1dn) lentiviral construct. j) Fragment length analyses of the three stable microsatellite loci in WT and BMRZTP organoids. Dashed black lines show average size of the WT loci. k) Selection of *Tgfbr2* knock-out (KO) in BMRZ organoids (BMRZT). In addition to TGFβ treatment, Noggin was withdrawn from the culture medium. Representative images after two passages of selection are shown. l) Nutlin-3a selection of *Tp53* ko. BMRZT organoids (BMRZTP). Representative images after two passages of selection are shown. m) Sanger sequencing results of the targeted regions in *Tp53* and *Tgfbr2*. At least 28 colonies were checked per gene and percentage of mutated sequences with or without frameshift (FS) are shown. Panels below show representative images of the loci targeted by the different sgRNAs. n) Volcanoplots displaying differential gene expression between BMRZTP vs. WT_wren and AKPS organoids vs. WT_wren. Threshold for significance in both was set at log2FC > 0.5, FDR < 0.05. Selected genes are labelled.

**Supplementary Figure 2.**
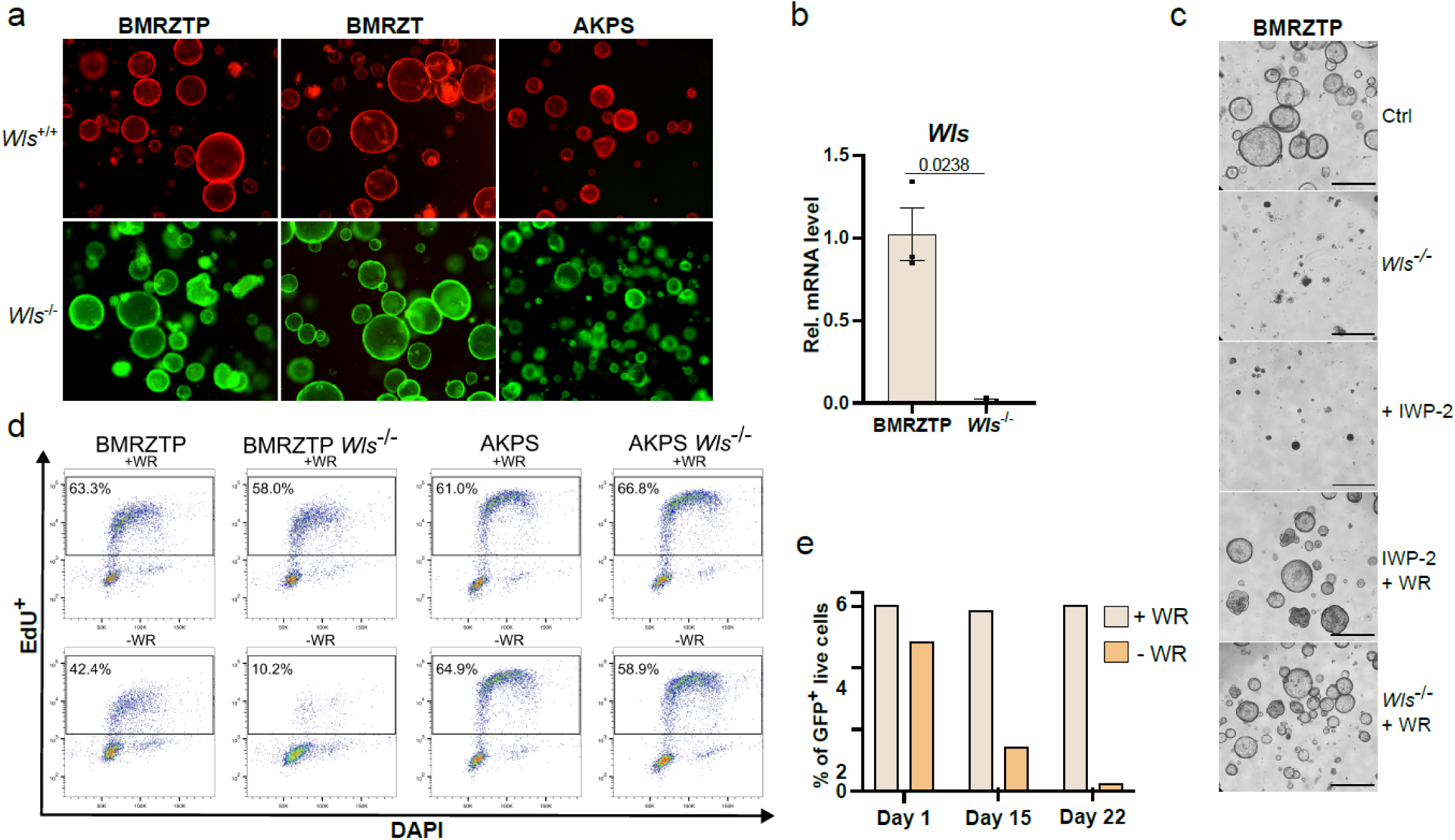
Testing WNT dependency in Apc- and Braf-driven CRC organoids. a) Representative images displaying TdTomato and GFP expression in sorted BMRZTP, BMRZT and AKPS organoids. TdTomato (magenta) indicates *Wls*^+/+^ cells, GFP (green) indicates *Wls*^−/−^cells. If not used for experimental purposes, *Wls*^−/−^organoids were routinely cultured in +WR medium. b) Relative *Wls* mRNA levels determined by RT-qPCR in BMRZTP and BMRZTP *Wls*^−/−^organoids. P-value was calculated by unpaired Welch’s test. c) WNT dependency tests in BMRZTP organoids. WNT secretion was either blocked by *Wls* elimination or treatment with the Porcn inhibitor IWP-2. Phenotypes were rescued by concomitant WR supplementation. Representative images after two passages are shown. Scalebar = 1mm. d) Representative FACS dot plots of the quantification shown in Fig. 2e. Black boxes represent the gates used to determine EdU positive cells. e) Quantification of the FACS dot plots of the time course experiment shown in Fig.2g. Bars represent the percentage of live GFP^+^ cells in chimeric BMRZT organoids in + WR or −WR medium over time.

**Supplementary Figure 3.**
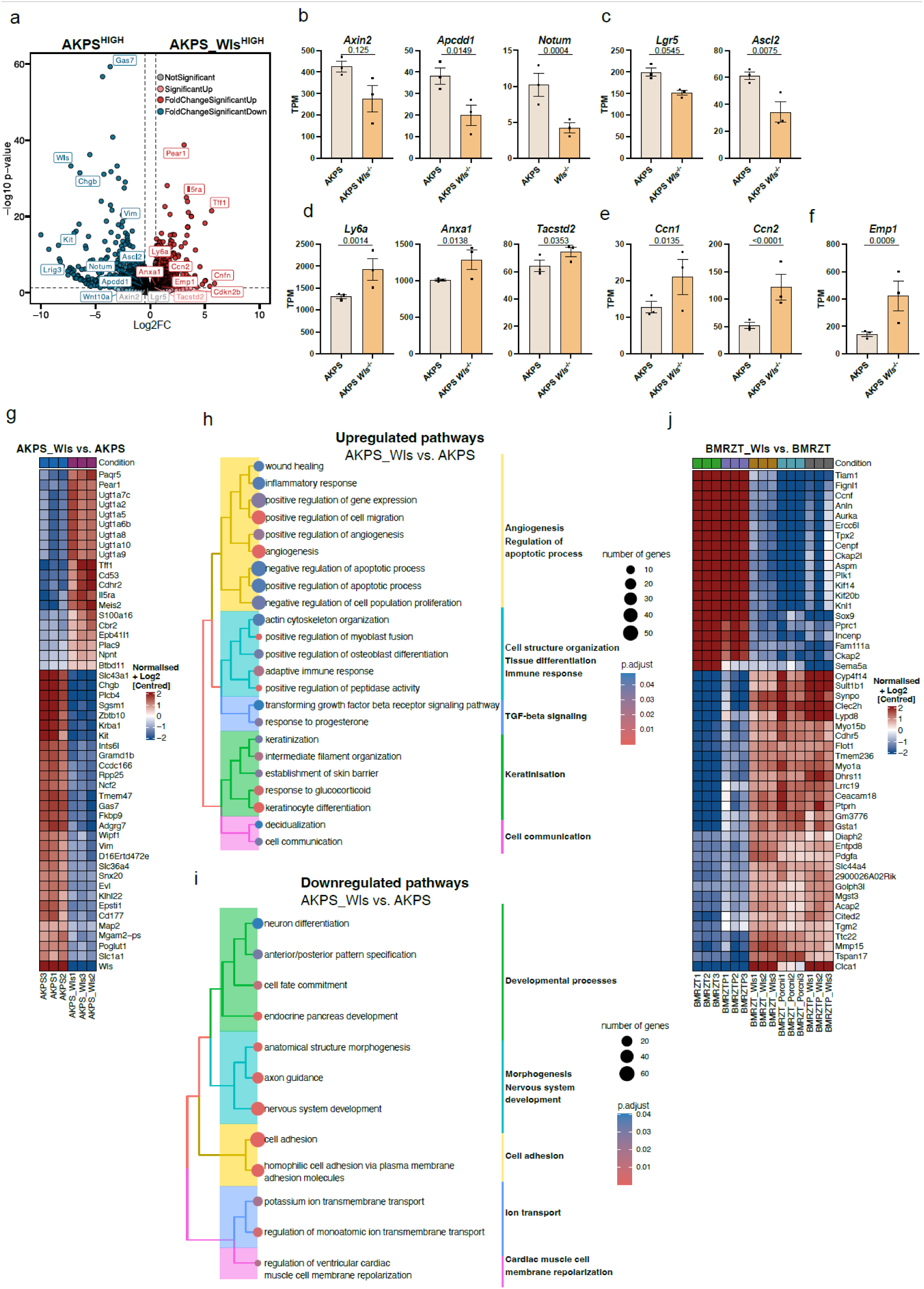
AKPS organoids remain responsive to loss of WNT secretion. a) Volcano plot showcasing transcriptomic differences between AKPS_Wls and AKPS organoids. Threshold for significance was set at log2FC > 0.5, FDR < 0.05. Selected genes are labelled. b) mRNA Expression levels of selected WNT target genes in TPM. FDRs were derived from DESeq2. c) Expression levels of colonic stem cell markers. d) Expression levels of fetal reprogramming marker genes. e) Yap target gene expression levels. f) Expression levels of *Emp1*. FDR values in b), c), d), e) and f) were calculated by DESeq2. g) Heatmap showing expression counts of the top 50 most differentially expressed features in AKPS_Wls versus AKPS by p-value. Both up- and downregulated DEGs are shown. Normalized, centered log2 counts are shown. Red color indicates upregulation, blue color indicates downregulation. h-i) ORA analysis of the top 30 up- and downregulated pathways between AKPS_Wls and AKPS summarized in tree-plots. Results are based on the “Molecular function” gene ontology (GO). Number of genes per GO term are indicated by dot size. Pathway summary was added manually. Color indicates statistical significance of pathway enrichment. Adjusted p-values are shown. j) Heatmap showing expression counts of the top 50 most differentially expressed features in BMRZT_Wls versus BMRZT by p-value. Both up- and downregulated DEGs are shown. For comparison, also BMRZT treated with porcni (BMRZT_Porcni), BMRZTP and BMRZTP_Wls are shown. Red color indicates upregulation, blue color indicates downregulation.

**Supplementary Figure 4.**
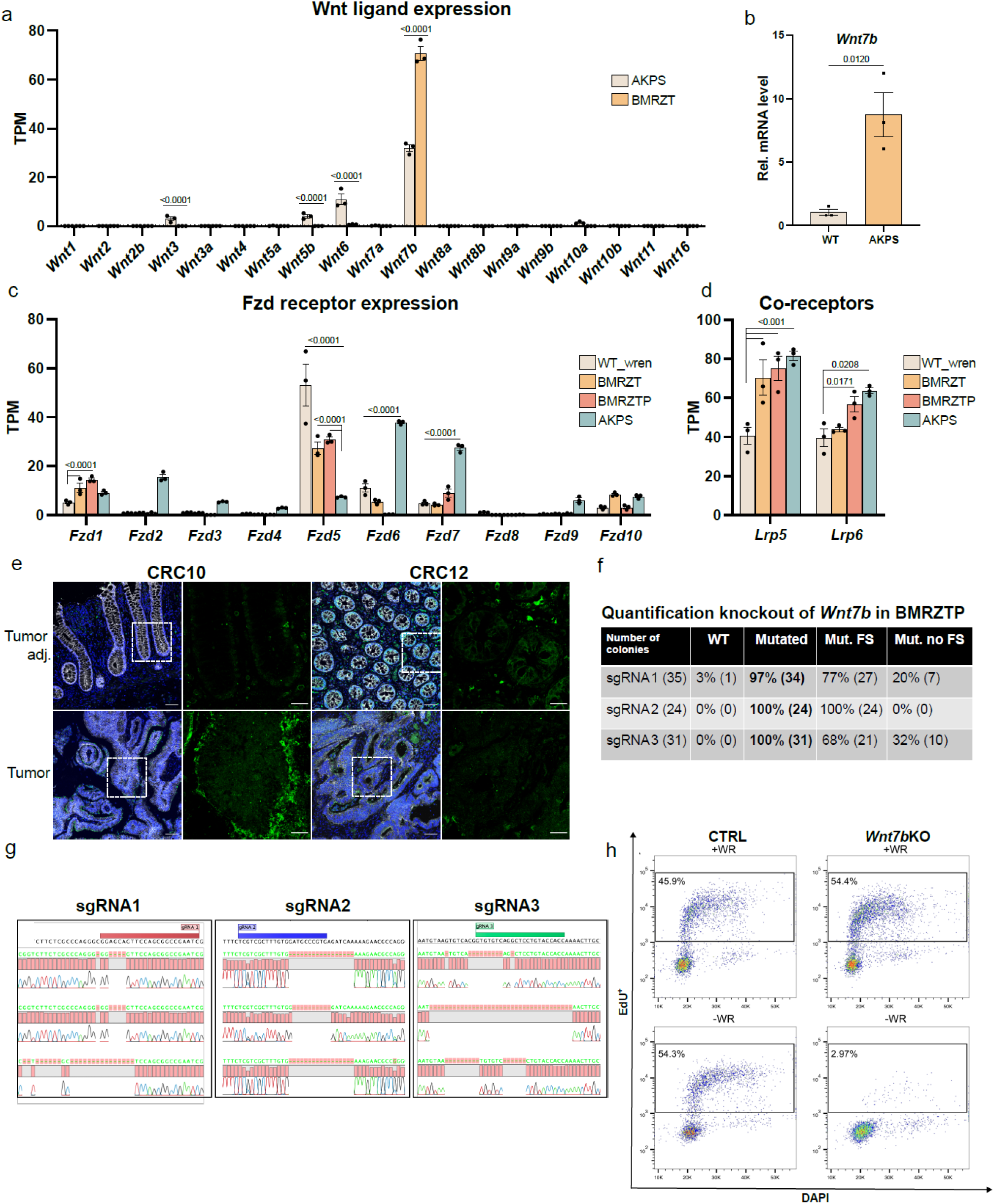
Validation of *Wnt7b* expression and knockout. a) Bar chart displaying TPM of all 19 WNT ligands in AKPS and BMRZT organoids. FDR values were calculated by DESeq2. b) RT-qPCR validation of *Wnt7b* expression in AKPS organoids. P-value was calculated by unpaired t-test. c) Bar chart displaying TPM from bulk of all 10 Frizzled (Fzd) receptors. FDR value was calculated by DESeq2. d) Bar chart displaying expression of Lrp5/6 co-receptors. FDR value was calculated by DESeq2. e) IF staining on human BRAF-mutant tumor and matched tumor-adjacent tissue. Staining shows WNT7B (green) E-cadherin (white) and DAPI (blue). Images were taken at 20x magnification with insets at 60x magnification (dashed white boxes). Scalebars: 60 and 25 μm. f) Sanger sequencing results of the three targeted *Wnt7b* exons. Per gRNA site, at least 31 colonies were analyzed. Percentage of mutated colonies with or without frameshift (FS) are shown. g) Panels show representative images of the sanger sequencing results at the distinct sgRNA binding sites in *Wnt7b* exons. h) Representative FACS dot plots of the EdU proliferation assay shown in Fig. 4h. Black boxes represent the gates used to determine EdU-positive cells.

**Supplementary Figure 5.**
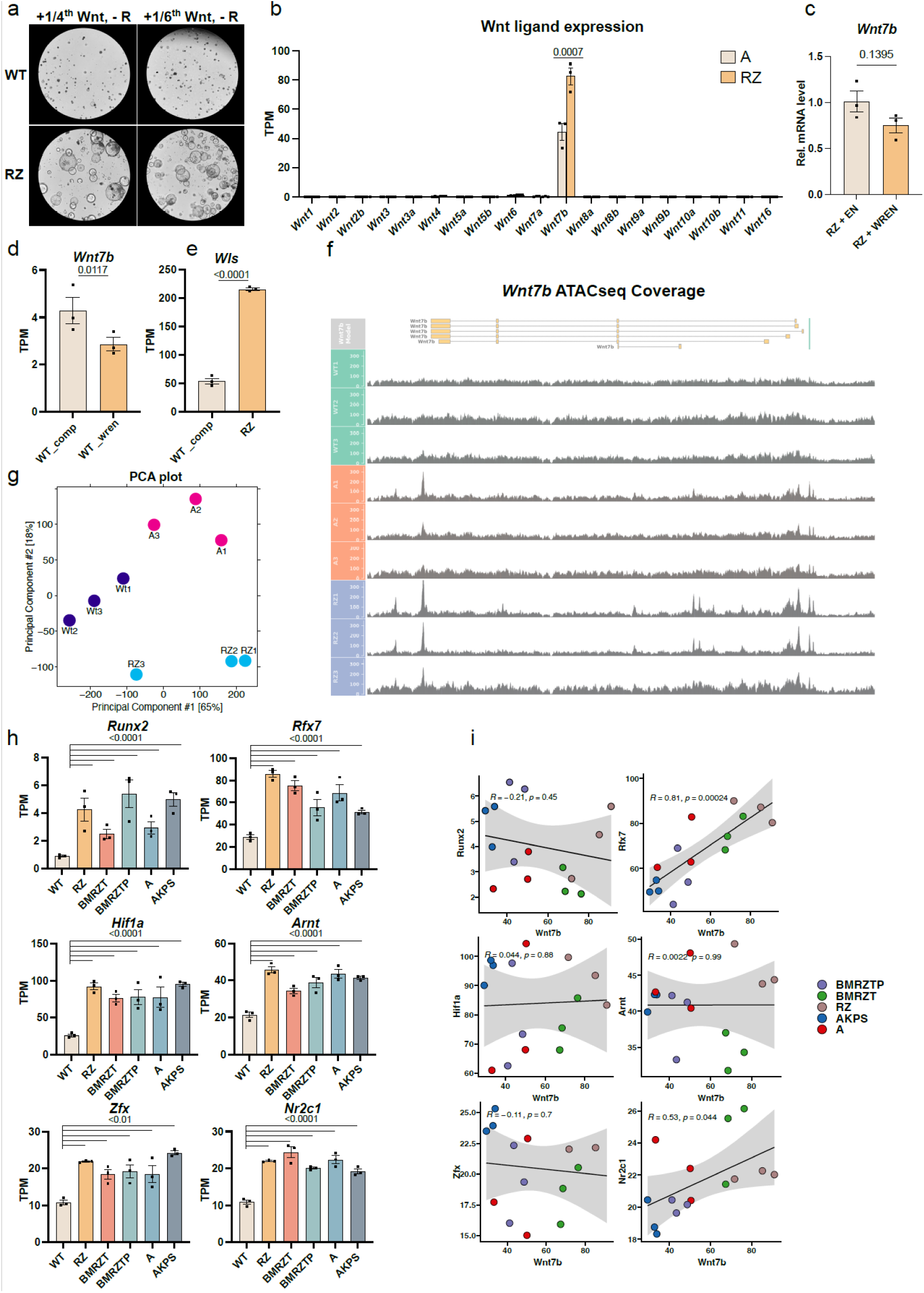
*Wnt7b* re-expression and potential candidates. a) Culture of RZ and WT organoids in decreasing concentrations of WNT surrogate. The standard concentration used was 0.5nM. 1/4^th^ WNT = 0.125nM, 1/6^th^ WNT = 0.033nM. b) Expression of all 19 WNT ligands measured as TPM in A and RZ organoids. Values for *Wnt7b* are also shown in Fig.5f. FDR value was calculated by DESeq2. c) Influence of WNT surrogate and Rspo (WR) supplementation on *Wnt7b* mRNA levels in RZ. Organoids were either supplemented with full medium (+WREN) or with EGF and Noggin (+EN). P-value was calculated using an unpaired t-test. d) Changes in *Wnt7b* levels upon WNT starvation in WT organoids. After passage, WT (Wt_comp) organoids were cultured in complete medium (+WREN). Wt_wren organoids were cultured one day in complete medium (+WREN) followed by two days in −WREN medium. FDR values were calculated by DESeq2. e) Increase in *Wls* mRNA in RZ vs Wt_comp organoids. FDR value was calculated by DESeq2. f) Overview of the chromatin accessibility at the *Wnt7b* locus. ATAC-seq peaks of the three replicates per condition are shown. Yellow boxes at the top indicate different *Wnt7b* transcript isoforms. g) ATAC-seq PCA plot of Wt_comp, A and RZ organoids. The three replicates per condition are shown. h) Expression profile of the different candidate TFs of *Wnt7b*. WT = Wt_comp. All candidates are differentially expressed compared to Wt_comp (log2FC > 0, FDR < 0.05). Expression levels are also shown in Fig.5h. i) Correlation of candidate TFs and *Wnt7b* expression in TPM. P-value was calculated using Pearson’s correlation test.

